# Cancer cells co-evolve with retrotransposons to mitigate viral mimicry

**DOI:** 10.1101/2023.05.19.541456

**Authors:** Siyu Sun, Jungeui Hong, Eunae You, Kaloyan M. Tsanov, Jonathan Chacon-Barahona, Andrea Di Gioacchino, David Hoyos, Hao Li, Hua Jiang, Han Ly, Sajid Marhon, Rajmohan Murali, Pharto Chanda, Ali Karacay, Nicolas Vabret, Daniel D. De Carvalho, John LaCava, Scott W. Lowe, David T. Ting, Christine A. Iacobuzio-Donahue, Alexander Solovyov, Benjamin D. Greenbaum

## Abstract

Overexpression of repetitive elements is an emerging hallmark of human cancers^1^. Diverse repeats can mimic viruses by replicating within the cancer genome through retrotransposition, or presenting pathogen-associated molecular patterns (PAMPs) to the pattern recognition receptors (PRRs) of the innate immune system^2–5^. Yet, how specific repeats affect tumor evolution and shape the tumor immune microenvironment (TME) in a pro- or anti-tumorigenic manner remains poorly defined. Here, we integrate whole genome and total transcriptome data from a unique autopsy cohort of multiregional samples collected in pancreatic ductal adenocarcinoma (PDAC) patients, into a comprehensive evolutionary analysis. We find that more recently evolved **S**hort **I**nterspersed **N**uclear **E**lements (SINE), a family of retrotransposable repeats, are more likely to form immunostimulatory double-strand RNAs (dsRNAs). Consequently, younger SINEs are strongly co-regulated with RIG-I like receptor associated type-I interferon genes but anti-correlated with pro-tumorigenic macrophage infiltration. We discover that immunostimulatory SINE expression in tumors is regulated by either **L**ong **I**nterspersed **N**uclear **E**lements 1 (LINE1/L1) mobility or ADAR1 activity in a *TP53* mutation dependent manner. Moreover, L1 retrotransposition activity tracks with tumor evolution and is associated with *TP53* mutation status. Altogether, our results suggest pancreatic tumors actively evolve to modulate immunogenic SINE stress and induce pro-tumorigenic inflammation. Our integrative, evolutionary analysis therefore illustrates, for the first time, how dark matter genomic repeats enable tumors to co-evolve with the TME by actively regulating viral mimicry to their selective advantage.

## MAIN

Repetitive elements account for approximately 54% of the human genome^6^, with more than half consisting of retrotransposable elements that can, when fully functional and activated, insert themselves into new genomic sites via a copy-and-paste mechanism^6^. SINE and LINE (**S**hort / **L**ong **I**nterspersed **N**uclear **E**lements) are two major classes of retrotransposable elements found in the human genome^6^. There are ∼500,000 copies of LINE-1 (or L1) occupying 21% of the human genome, but only ∼100 full-length human-specific L1 elements (L1Hs) are capable of active and autonomous retrotransposition^7^. L1-encoded proteins also mobilize non-autonomous retrotransposons – and these retrotransposed SINEs, make up another 13% of human DNA^1^. Due to the deleterious nature of retrotransposition, multiple defense mechanisms, including epigenetic silencing and post-transcriptional degradation, have evolved to protect genomes^8, 9^. These protective functions are often dysregulated in cancer^10^ or aging^11^, resulting in the frequent observation of derepressed repeats in multiple cancer types^12, 13^ and older individuals^11^. The dysregulation of SINEs and L1 can induce “viral mimicry” by enabling both reverse transcription and the display of pathogen-associated molecular patterns (PAMPs) which have the potential to trigger inflammatory responses^2–5^. Our work in ovarian and pancreatic cancer has found an enrichment of repeat RNAs in tumor-derived extracellular vesicles (EVs) capable of stimulating monocyte-derived macrophages, demonstrating these viral-like repeat sequences can, in principle, alter the tumor microenvironment (TME)^13^. However, the extent to which tumors use viral mimicry from retrotransposable elements to co-opt the immune system to their selective advantage remains unclear.

Pancreatic ductal adenocarcinoma (PDAC) is a devastating disease and the fourth leading cause of cancer-related deaths in the United States. Despite an increase in five-year patient survival from 8% to 11% during the past five years^14^, the disease remains largely incurable. Inflammatory processes have emerged as key mediators of pancreatic cancer development and progression, with inflammation associated with negative outcomes by forming a highly immune-suppressed TME^15, 16^, implying tumors modulate the immune system to their selective advantage^15, 16^. Like many other tumors, PDAC exhibits abnormal expression of repetitive elements, with high enrichment in the over 70% of cases with *TP53* mutations^12, 17^. We conducted an integrative genomic analysis of 214 multiregional rapid autopsy samples from PDAC patients, demonstrating for the first time the co-evolution of mutant *TP53*, inflammation, retrotransposition and the display of viral mimicry in PDAC. As validated in an additional large cohort of PDAC tumors and analysis of PDAC cell lines, we found immunostimulatory SINE expression is negatively regulated by L1 retrotransposon activity in the setting of *TP53* mutation while, in the setting of wildtype *TP53* status, tumors utilize ADAR1 as a response mechanism to SINEs. Both mechanisms highlight the multiple pathways tumor cells utilize to converge on repeat regulation to promote tumor growth via immune manipulation.

### SINE elements are co-expressed with RIG-I like receptor (RLR) associated interferon signaling in PDAC autopsy cohort

We integrated multi-omics data, including whole genome and total RNA sequencing collected from a rapid autopsy PDAC cohort^18^ to characterize the role of repetitive elements in mediating tumor inflammation and evolution (**Fig. 1A**). To our knowledge, this is the largest group of total RNA sequencing human specimens in PDAC, a sequencing protocol necessary for unbiased assessment of repeats^18^. Our cohort also covers a substantial number of samples from primary and metastatic sites from multiple tissues, allowing us to track tumor evolution within each patient. In total, whole transcriptome data was collected from 214 multiregional samples across 27 patients (**Fig. 1B**). Additionally, 58 matched tumor samples from 8 patients were simultaneously subjected to whole genome sequencing (WGS) (**Fig. 1B**), allowing us to, for the first time, disentangle the interaction between repeat transcription, innate immunity, and tumor genome evolution.

**Figure 1.**
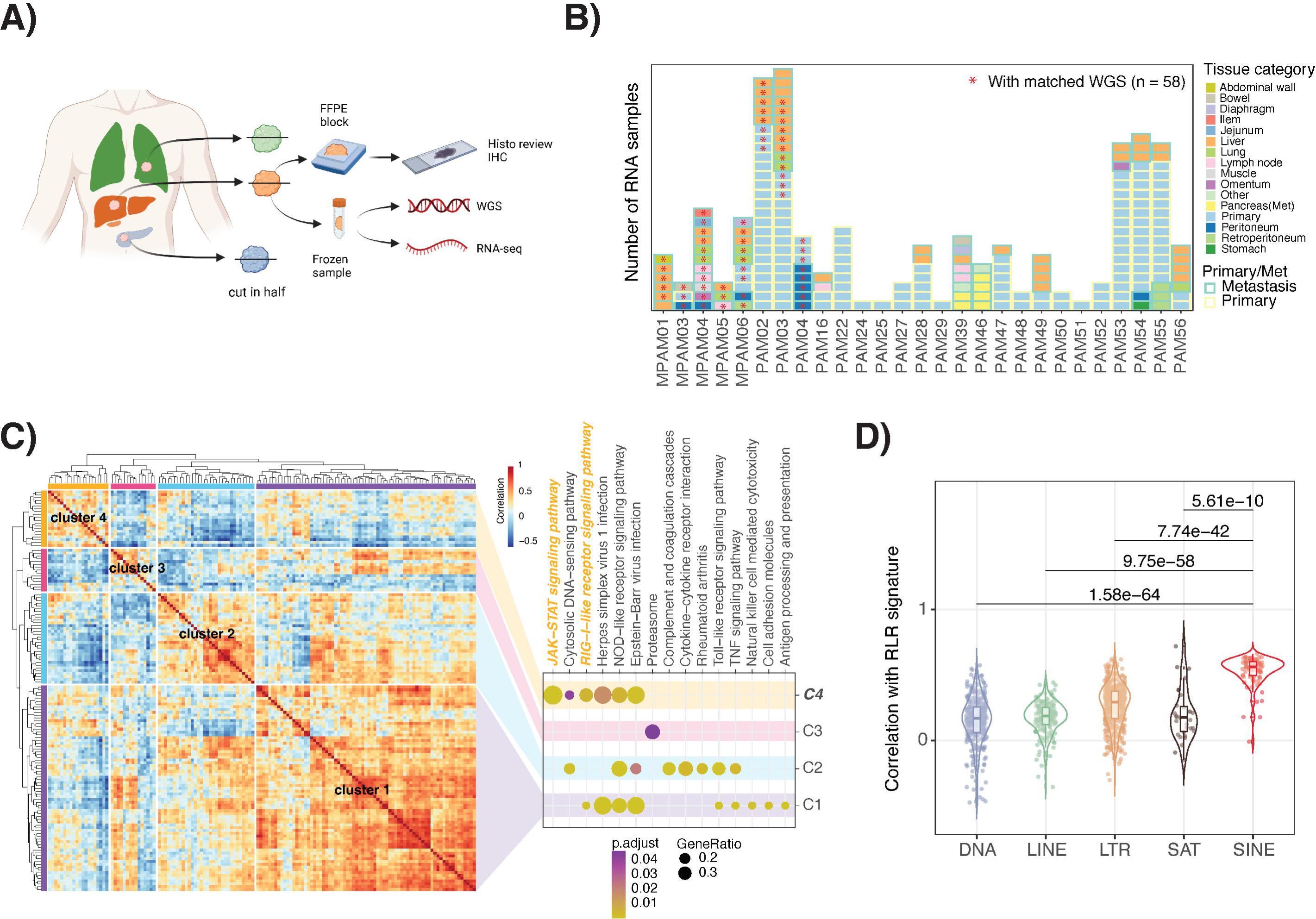
SINE elements are co-expressed with RIG-I like receptor associated IFN signaling in PDAC autopsy cohort. **A)** Multiregional samples and multi-omics data overview of autopsy PDAC cohort. **B)** Sample composition of the autopsy PDAC cohort: 214 total RNAseq samples with 58 of which have matched WGS. **C)** Pairwise correlation analysis of 140 interferon related genes expression in 214 samples revealed 4 co-expression clusters (left) (**Table S1**). KEGG pathway annotation of genes identified distinct functional pathways in each cluster (right). Cluster 4 (C4) exclusive pathways are highlighted in yellow. **D)** Pearson correlation coefficient between median expression of genes in C4 (hereafter RLR signature) and repeat expression in different classes across 214 samples. Benjamini-Hochberg corrected P values of t-test are labeled.

To assess the relationship between repeats and innate immune signaling, we quantified repeat expression^19^, and evaluated differentially expressed genes and repeats in tumors collected from primary versus metastatic sites. Given the relatively high stromal content in PDAC even after laser microdissection, we performed tumor purity corrected differential expression analyses (**Methods, Fig. S1A**)^20^. Metastatic tumors were found to be enriched for multiple inflammatory signatures using Gene Set Enrichment Analysis^21^ compared to primary tumors. Meanwhile, five immune related HALLMARK pathways including interferon alpha and gamma responses were upregulated in metastatic samples compared to primary tumors (**Fig. S1B**), suggesting a selective advantage for inflammatory states in metastasis.

We then focused on the type I interferon pathways which have shown associations with repeat-mediated viral mimicry in multiple cancer types^5, 22^. We curated a list of 140 interferon genes (IFN) and interferon stimulated genes (ISG) (**Table S1**). Using unsupervised clustering on gene expression, we identified four co-expression clusters out of the 140 genes using pairwise correlation across all 214 samples (**Fig. 1C**). Further KEGG pathway annotation of genes identified distinct functional pathways enriched in each cluster. Notably, retinoic acid–inducible gene I (**R**IG-**I**) **L**ike **R**eceptor (RLRs) pathway and JAK-STAT signaling pathway are over-represented in one cluster – labelled C4. Recent studies have revealed that both viral and host-derived RNAs can trigger RLR activation^23–25^, we therefore sought to identify the association between expression of specific repeat classes (SINE, LINE, LTR/ERVs, SAT, DNA) and different clusters. Among five repeat classes, SINEs show the most significant and strongest positive correlation with the RLR signature (median r = 0.53) (**Figs. 1D, S1C**), which is not affected by the absolute expression level of SINEs (**Fig. S1D**). The association between highly coregulated core RLR mediated IFN signatures and SINEs expression, supports the notion that SINEs^26^, and not **E**ndogenous **R**etro**V**irus (ERVs/LTRs)^5^, are likely the primary binding substrate of RLRs in PDAC. Our result is in line with a recent study which claims that a subtype of SINE – inverted Alu elements are the major substrates of MDA5, a known RLR *in vitro*^3, 4^.

### Newly evolved SINEs show stronger correlation with RLRs signature and are prone to form dsRNA

To dissect which SINE subfamilies are responsible for the strong interaction with RLR mediated signaling, we stratified SINEs into six groups based on their evolutionary age (**Fig. 2A, Methods**)^27–32^. The positive correlation between RLR associated IFN signature and each SINE subgroup increased as the evolutionary age decreased, with AluY harboring the largest variations (**Fig. 2A**). Moreover, unbiased clustering of all genes defining the RLR signature, and all SINE elements revealed two gene/repeat clusters, where younger SINEs are grouped together with 17/20 genes in the RLR signature (**Fig. S2A**). Altogether those results imply that younger SINEs have a stronger association with RLR signature in PDAC.

**Figure 2.**
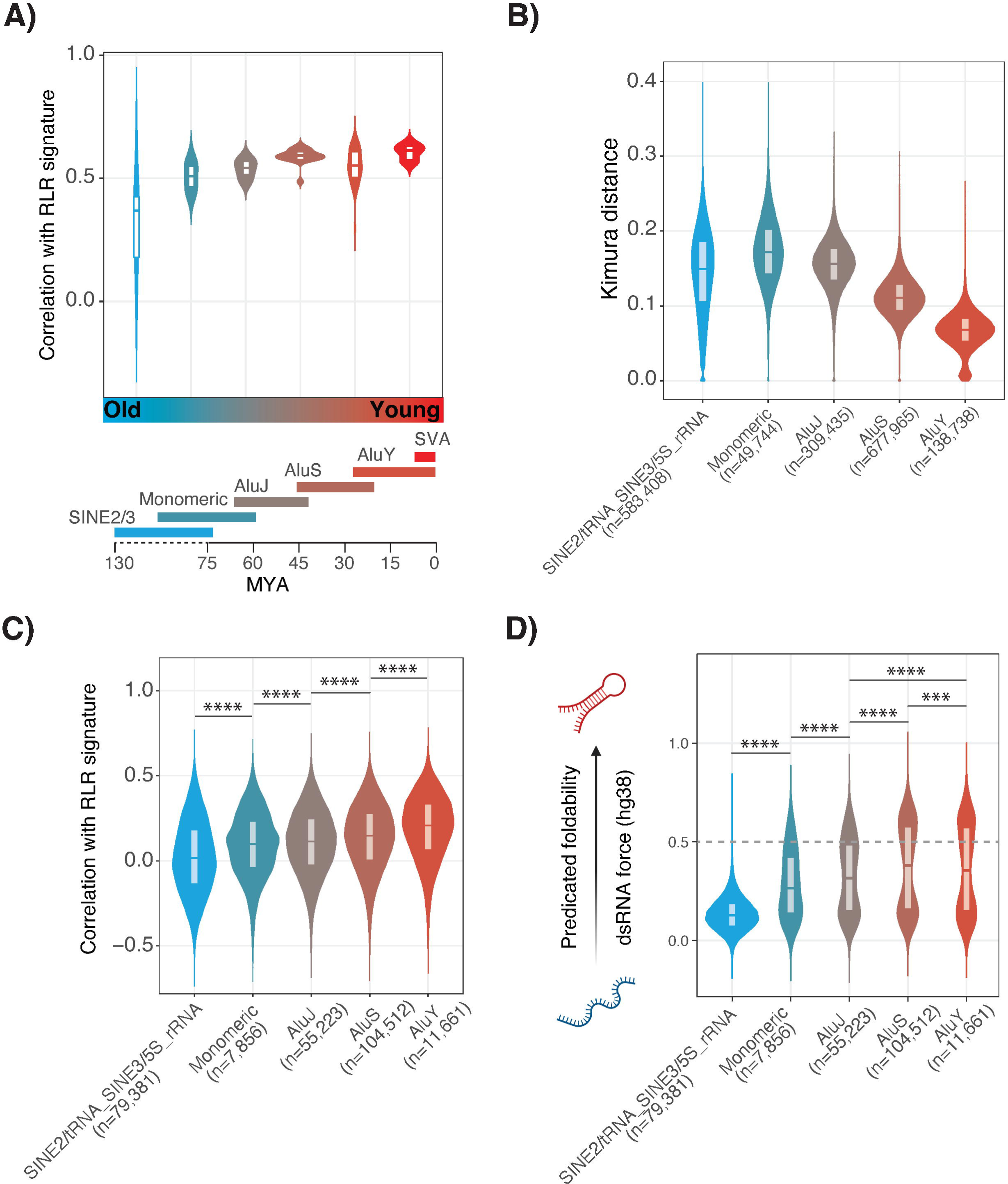
Younger SINE elements have stronger correlation with core RLR signature and are prone to form dsRNA. **A)** Pearson correlation between RLR signature and different SINE subtypes. SINE subtypes are colored based on their evolutionary age. Evolutionary timeline of SINEs: SINE2/SINE3, SINE1 (Monomeric and dimeric AluJ, AluS and AluY, and SVA). **B)** Kimura distance of all copies of SINEs from the consensus sequence of the subfamily grouped by SINE evolutionary groups. **C)** Pearson correlation coefficients between RLR signature and locus specific expression of SINEs stratified by evolutionary age groups. **D)** dsRNA forces calculated for the human genome (hg38) that encompassing SINEs. (* p < 0.05, ** p < 0.01, *** p< 0.001, **** p < 0.0001, t-test).

We then investigated the relation between immunogenicity and evolutionary age at the level of single copies. We quantified the evolutionary age of each SINE copy by utilizing the evolutionary distance of each copy against its reference consensus sequence^25^ (**Fig. 2B**). Consistent with the general association of younger SINEs and RLRs, the association with the RLR signature is significantly higher when SINEs are evolutionarily younger (**Fig. 2C**). Given that RLRs typically recognize and bind to double-stranded RNA (dsRNA) to initiate a downstream innate immune response^33^, we hypothesized that younger SINEs possess a greater potential to generate dsRNAs than older SINEs. To test this hypothesis, we adapted our previously developed computational framework^25^ for assessing the potential of a specific SINE copy to fold into a dsRNA with a nearby sequence (< 3000bp), which we referred to as the dsRNA force (**Methods**). SVAs, as a composite SINE, are excluded from this analysis as their sequence composition and size is quite different from other SINEs. The distribution of dsRNA forces is also evolutionary age depended, with younger groups - AluY and AluS harboring significantly higher dsRNA force than older groups (**Fig. 2D**). However, AluS has a slightly higher fraction of dsRNA forming copies compared to AluY (**Fig. S2B**), likely indicating a secondary selection pressure on AluS in maintaining their capability to form dsRNA. SINEs with stronger predicated foldability (dsRNA force > 0.5) have significant stronger association with RLR signature (**Fig. S2C**). Moreover, the mean foldability of each SINE subfamily significantly correlated with the association with pre-defined core IFN signatures (r = 0.77, p < 0.00001) (**Fig. S2D**). Taken together, our findings suggest that the expression of younger SINEs is more strongly linked to RLR-mediated innate immune responses due to their increased capacity to form dsRNAs.

### L1 transposition frequency is associated with mutant *TP53* status and continues throughout tumor evolution

L1 is the sole active autonomous retrotransposable element in the human genome, and its activity has the potential to influence the genomic evolution of cancer cells^34^. To precisely identify L1 insertions, we implemented a pipeline^35^ that utilizes soft-clipped reads and discordant read pairs derived from non-reference L1 insertion sites present only in tumor samples compared to matched normal samples. In total, we identified 373 *de novo* L1 insertions in 39 WGS samples (**Table S2**) (out of 74) from five patients (out of eight), with a substantially greater number of insertions identified in samples obtained from metastatic sites (**Fig. 3A**). Out of the 39 samples with detected *de novo* L1 insertions, only one was obtained from the primary site. No common insertions were detected across patients and the insertion sites overlap across different metastatic sites or patients were limited (**Fig. 3B**). One patient (MPAM05) exhibited nearly a tenfold increase in the number of *de novo* L1 insertions compared to the other patients (**Fig. 3A**), with exclusive nuclear L1 ORF1p localization as assessed by immunohistochemistry (**Fig. S3A**). Consistent with the result from an orthogonal quantification method for L1 retrotransposition^36^, the expression level of the L1 ORF1p, as scored by immunohistochemistry, correlated with the number of *de novo* insertions (r = 0.48), albeit not to a statistically significant degree (p = 0.054) (**Fig. S3B**). Different from patient MPAM05, IHC of L1 ORF1p from the other patients show a previously documented cytoplasmic localization (**Fig. S3B**).

**Figure 3.**
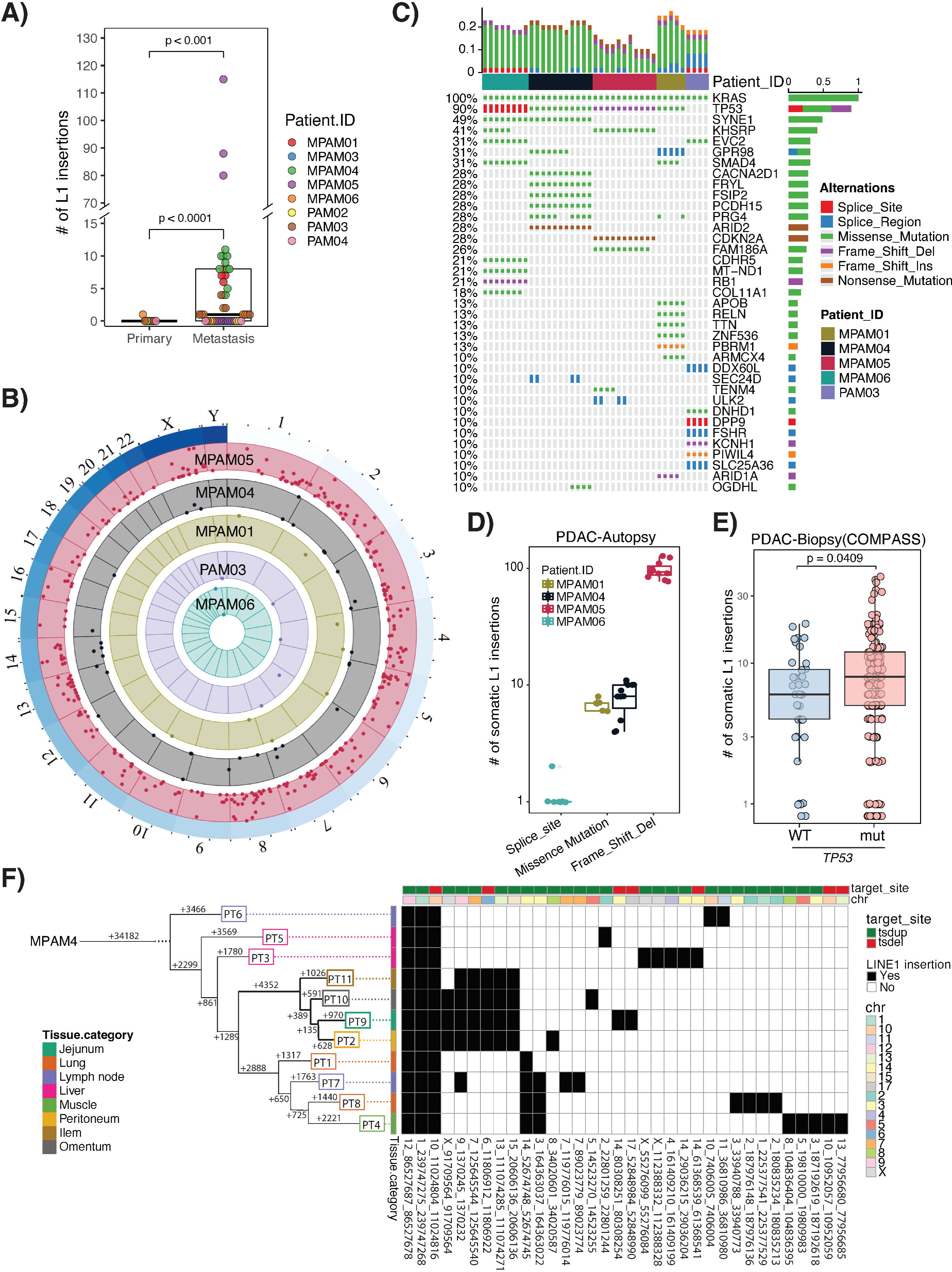
L1 retrotransposition activity is associated with mutant *TP53* status and tracked with phylogenetic tumor evolutionary branching. **A)** Overview of *de novo* tumor specific insertions of LINE1 in 74 samples from 8 patients based on Whole Genome Sequencing (WGS) data stratified by tumor sites (primary vs metastasis). Out of which, 31 samples from 5 patients have non-zero somatic L1 insertion. **B)** Insertion loci overview of 31 samples from 5 patients have non-zero somatic insertion detected. Track background is colored by the patient. Each dot represents one insertion locus that past QC (**Methods**). Position of each dot is proportion to number of samples share the same insertion from the same patient. For example, dots on the inner margin of each patient indicate the insertion is only found in one sample, dots on the outer margin indicate that insertion happened in all samples for that given patient. **C)** Oncoprint of SNPs with frequency over 10% across samples from the 5 patients. All three samples from patient MPAM05 have a frameshift deletion in *TP53* at c.78delT. **D)** Boxplot of number of L1 insertion in samples with different *TP53* mutation status in PDAC autopsy cohort. Multiple dots per box indicate multiregional samples from the same patient. Given that the samples in each *TP53* mutation background are not independent, no statistical test were applied. **E)** Boxplot of number of somatic L1 insertion in samples with wild type or *TP53* mutants in PDAC biopsy cohort in COMPASS trial^36^. p-value representing the t-test significance of somatic L1 insertion difference between WT and *TP53* mutant samples is labeled. **F)** Integrative analysis of L1 insertion phenotypes and tumor evolution of patient MPAM04. Samples are boxed with colors representing the tissue type. Target site features is coded as red (target site duplication) and green (target site deletion).

Subsequently, we evaluated potential genetic factors that may account for the significantly higher number of L1 insertions observed in patient MPAM05. No significant association was detected between the number of L1 insertions and either the COSMIC mutational signature or microsatellite instability. Notably, an analysis of SNP calls on the WGS data revealed a distinct frameshift deletion of *TP53* in all 11 samples obtained from MPAM05 (**Fig. 3C**). The specific frameshift deletion eliminates the p53 DNA binding domain, resulting in a complete loss of function for the p53 protein. Upon comparing the frequency of L1 insertions across the 39 samples stratified by different *TP53* mutation variants, we observed an association between L1 retrotransposition frequency and their *TP53* mutation status (**Fig. 3D**). Samples with splice site mutations (MPAM06) had the lowest insertion frequency, which is comparable to patient PAM03 who has a wildtype *TP53*. However, samples with missense mutations (MPAM01& MPAM04) had a higher number of insertions. The highest insertion frequency was observed in samples with frameshift deletions – MPAM05 (**Fig. 3D**). To further validate our findings, we conducted the same analysis on WGS from an independent biopsy cohort from the COMPASS trial^37^. Consistently, metastatic samples had a higher number of somatic L1 insertions compared to primary samples (**Fig. S3C**). Furthermore, samples harboring *TP53* mutations tend to have a higher number of insertions than those without mutations, with higher number of insertions in los of function *TP53* mutation types (**Figs. 3E, S3D, E**). Those results have fortified the role of p53 as surveyor of L1 mobility in PDAC, also corroborated in other different cancer types^35, 37^.

To investigate the L1 re-integration trajectory along tumor evolution, we utilized WGS data to reconstruct a phylogenetic tree of metastases and subclones based on their anatomical locations^39^. In patient MPAM04, all samples had three *de novo* insertions in common on chromosomes 1, 10, and 12. Samples within the same clade shared common insertions, as seen with samples PT2, PT9, PT10, PT11, which all had 6 common insertions located on different chromosomes (**Fig. 3F**). We observed similar concurrency between tumor evolution and L1 insertion in patient MPAM05 (**Fig. S3F**), where samples that are evolutionarily close share similar insertions. However, due to the limited samples, we were not able to perform a similar comparison for samples from the other three patients. The phylogenetic relationships of L1 insertions were not confined to a shared anatomic location or tissue type, suggesting that L1 activity does not follow a linear evolution along tumor progression. Instead, L1 insertions seem to be seeded by multiple prior locations and then disseminated to different anatomic locations. These results indicate that the majority of L1 insertions happened during the metastasis, which suggests a closer connection between L1 activity and tumor progression (e.g. metastasis/tumor dissemination), rather than just during tumorigenesis (e.g. initiation/local progression).

### Immunogenic SINE expression is anti-correlated with L1 retrotransposition and M2 macrophage infiltration

L1 retrotransposition is dependent on the expression of L1 proteins, such as ORF1p and ORF2p, as reintegration of L1 requires reverse transcription of L1 mRNA by L1 proteins binding *in cis*. In addition, SINEs can utilize L1 proteins *in trans* to facilitate their own retrotransposition back into the genome^40^. Hence, we next explored the interplay between L1 retrotransposition, SINE expression, and the core RLR signatures linked to younger SINE elements. SINEs exhibited a unique pattern compared to other repeat classes (LINE, ERV/LTR, DNA, SAT) and were consistently downregulated in samples with high L1 retrotransposition (**Fig. 4A**). More importantly, the antagonistic relationship between L1 mobility and SINEs was dependent on evolutionary age, with younger SINEs (SVA, AluY, AluS) having a stronger inverse relationship (**Fig 4B**). Including the three samples from patient MPAM05 did not change the observed trends (**Fig. S4A**), indicating our results are not biased by the presence of this patient’s high-integration samples. Consistently, the RLR-associated genes defined in **Fig 1C** are also repressed in samples with high number of L1 insertions (**Fig. S4B**). In addition, SINEs showed the strongest negative ranked correlation with the number of L1 insertion compared to other types of repeats (**Fig. S4C**), and the evolutionary younger SINEs are negatively associated with L1 activity, while older SINEs mostly show positive or no strong association (**Fig. S4D,** blue/grey vs red dots). These results suggest increased L1 retrotransposition may be selected for during tumor evolution due to its role in suppressing the immune response stimulated by dsRNA-forming SINEs.

**Figure 4.**
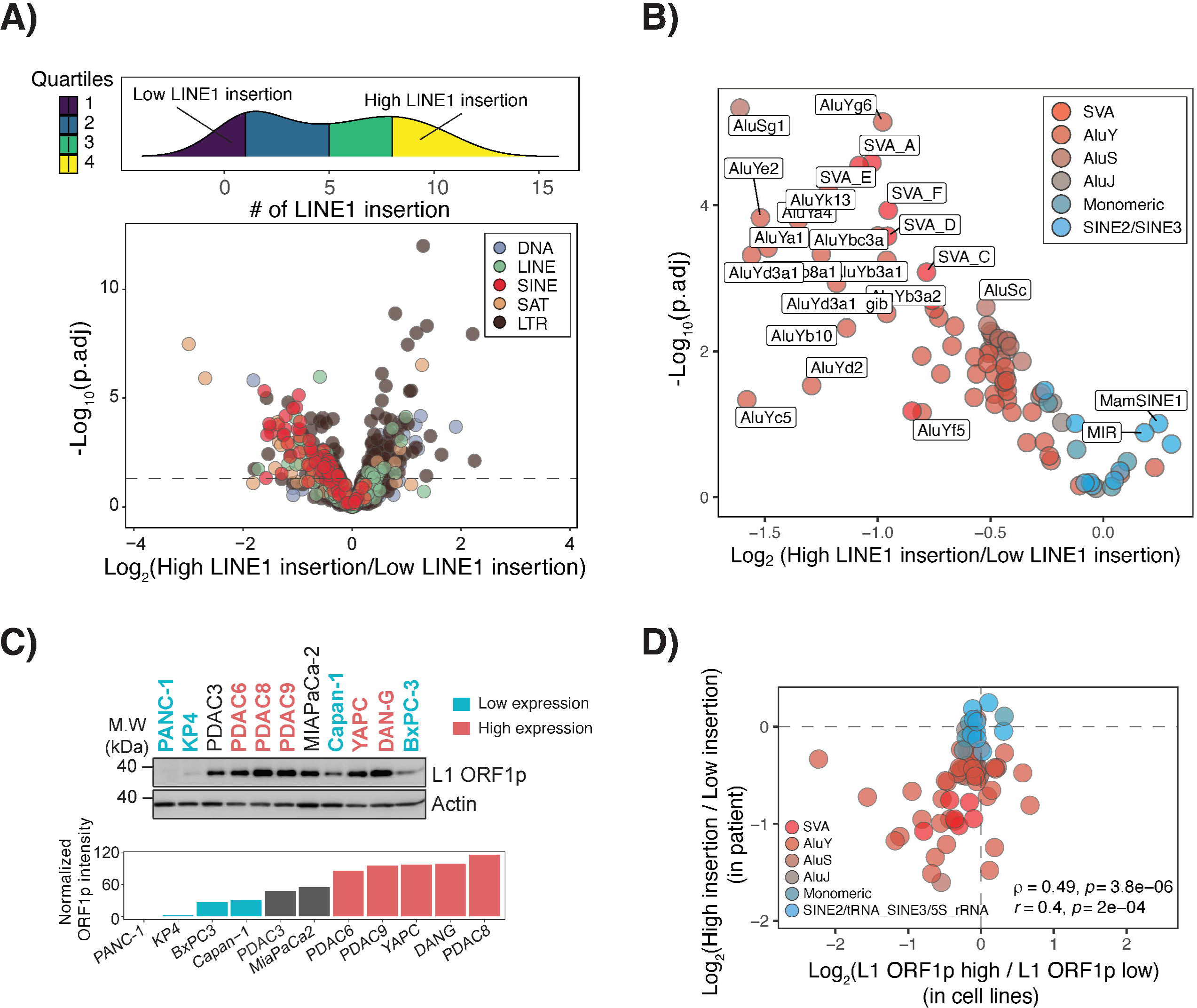
Immunogenic SINE expression is anti-correlated with L1 retrotransposition. **A)** Density plots colored by quartiles of samples (n=27, excluding 3 samples from MPAM05 with an extremely high number of somatic L1 insertions) with different number of somatic L1 insertions. Samples in quartile 4 (Q4) and quartile 1 (Q1) are selected to represent L1 insertion high and low samples for differential expression analysis shown as volcano plot below. The calculated Log_2_FoldChanges have been corrected for tumor purity (**Methods**). **B)** Volcano plot showing the differential expression of SINEs between samples with high (Q4) or low number (Q1) of somatic L1 insertions. SINE subtypes are colored based on their evolutionary age. **C)** Cropped western blot of L1 ORF1p expression in 11 PDAC patient derived cell lines. Actin normalized L1 ORF1p intensity were ordered to select for the high and low L1 ORF1p expression cell lines (**Table S3**). **D)** Comparison of change of SINEs expression between PDAC patient samples (Log_2_FC (high L1 insertions/low L1 insertions)) and *in vitro* cell lines Log_2_FC (high ORF1p expression/low ORF1p expression).

Next, we examined the relationship between the expression of different repeat classes and their associated immune infiltration. We utilized CIBERSORTx to perform cell type deconvolution from bulk RNA sequencing data. Out of all myeloid cells, M2 macrophages show a clear separation among different repeat classes (**Fig S4E**), a distinct evolutionary age dependent negative correlation between SINE expression and M2 macrophage infiltration (**Fig. S4F**), and subsequently a positive correlation between number of somatic L1 insertions and M2 macrophage infiltration (**Fig. S5A**). We attempted to validate the observed correlation between the expression of immunogenic SINEs, L1 retrotransposition, and immune infiltration in the COMPASS trial. However, due to the implementation of polyA selection in the COMAPSS trial RNA sequencing protocol, the significantly decreased ratio of reads mapped to repeats and coding genes prevent a similar comparison (**Fig. S5B**)^19^.

To corroborate the observed antagonistic relationship between L1 retrotransposon activity and SINE expression, we examined SINE expression in 11 PDAC cell lines with varying levels of L1 ORF1p expression quantified by western blot, as a proxy for L1 mobility (**Fig. 4C**). We then performed differential expression analysis between cell lines with low L1 ORF1p expression (PANC-1, KP-4, BxPC-3), and high L1 ORF1p expression (PDAC8, DAN-G, YAPC) (**Table S3**). We observed a consistent evolutionary age dependent downregulation of SINEs in samples with high L1 ORF1p expression (**Fig. S5C**). Together, our findings revealed a strong negative correlation between younger SINEs and L1 activity (**Fig. 4D**). Notably, SINEs demonstrated the highest level of consistency among repeat classes in the differential expression analysis in PDAC cell lines and PDAC human samples when stratified by L1 expression or activities (**Fig. S5D**), indicating a strong association between SINEs and L1 compared to other repeat classes.

### L1 ORF1p suppresses immunogenic SINE expression *in vitro*

To validate the direct repression of immunogenic SINE RNA by L1 ORF1p, we designed and tested two short hairpin (sh)RNA oligos (**Fig. 5A**) to knock down L1 ORF1p expression in PDAC6 cell line, which has relatively high L1 ORF1p expression (**Fig. 4C**). The knockdown experiment was performed in biological triplicates, and different knockdown efficiencies were observed among biological replicates and two shRNA oligos as revealed by western blot (**Fig. 5A**). We used the resulting distribution of samples with different L1 ORF1p expression levels to assess the impact of L1 ORF1p knockdown on repeat expression, calculating the correlation between GAPDH-normalized ORF1p intensity and repeat expression across all nine samples (**Fig. 5B**). In line with previous observations in the autopsy PDAC cohort and PDAC cell lines, SINEs showed the strongest negative correlation with L1 ORF1p protein expression (**Fig. S6A**), with younger SINEs being more prone to be repressed under high L1 expression (**Fig. S6B**). Collectively, SINEs constitute half and of the repeats that are significantly and negatively correlated with L1 ORF1p expression, while LINEs and LTRs were the only two classes of repats that showed significantly positive correlation with L1 ORF1p expression (**Fig. 5C**).

**Figure 5.**
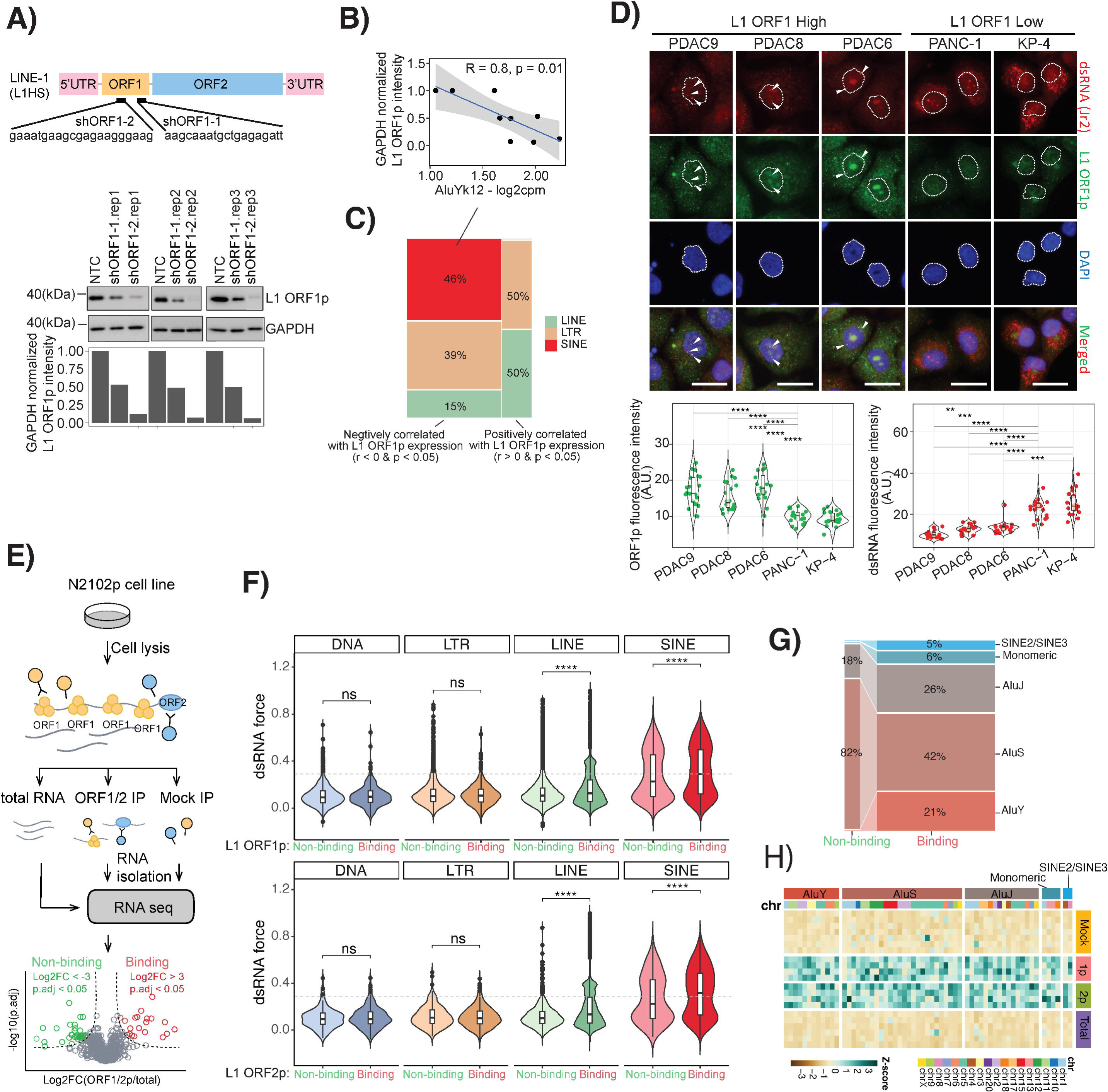
L1 retrotransposition activity suppresses immunogenic SINE expression likely via *in-cis* binding. **A)** Design of two short hairpin (sh)DNA sequences for knocking down of L1 ORF1p protein, and results expression of L1 ORF1p confirmed by western blot. GAPDH normalized ORF1p intensity were quantified. The knockdown experiment was performed in biological triplicates. **B)** An example scatter plot showing the anti-association between L1 ORF1p expression and young SINE (AluYk12) expression. **C)** Summarized compartmentalization of repeats that show significant negative (r < 0 & p < 0.05) or positive (r > 0 & p < 0.05) correlation with L1 ORF1p expression in the L1 shDNA knocking down experiments. SINEs are exclusive and dominating the repeats that are negatively associated with L1 ORF1p expression. **D)** Immunofluorescent staining with dsRNA antibody (Jr2), L1 ORF1p antibody, and DAPI in L1 ORF1p high (PDAC6, 8, 9) and L1 ORF1p low (PANC-1, KP-4) cell lines (**Methods**). Intensities of L1 ORF1p green signal and dsRNA red signal are quantified for each cell line and plotted on the right violin plots. **E) R**NA **I**mmuno**P**recipitation **seq**uencing (RIPseq) experimental setup and computational pipeline for detecting ORF1p and ORF2p binding transcripts in embryonal carcinoma cell line – N2102p. **F)** Violin plots show the distribution of dsRNA force assigned for each repeat copy (**Methods**) grouped by whether it is bound by protein of interests – ORF1p and ORF2p detected in the RIPseq. p-values calculated by t-test are labeled (**** p.value < 0.00001). **G)** Composition of SINEs bound by both L1 ORF1p and ORF2p. SINE subtypes are colored based on their evolutionary age. **H)** Heatmap shows the expression (Z-score) of L1 ORF1p and ORF2p binding SINEs in the RIPseq experiment.

We then sought to verify whether the observed reduction of immunogenic SINE expression leads to a reduction in dsRNA production. We performed immunofluorescent staining with anti-dsRNA (rJ2 clone) antibody (**Methods**) on five selected cell lines, considering cytoplasmic volume for the further analysis. Representative immunofluorescence images of cytoplasmic dsRNA have shown a significant reduction in high L1 ORF1p expressing PDAC cell lines (PDAC6, PDAC8, PDAC9) compared to low L1 ORF1p cell lines (KP-4, PANC-1) (**Table S3**). We found that high L1 ORF1p expressing cell lines had a reduced signal from the dsRNA antibody rJ2 in immunofluorescent imaging analysis (**Fig. 5D**). Furthermore, we observed a co-localization phenotype of L1 ORF1p and dsRNA antibodies in the nucleus of cells carrying high L1 ORF1p expression. These findings provide further evidence for a potential link between L1 activity and dsRNA accumulation in cancer cells.

The observed co-localization of L1 ORF1p and dsRNA in PDAC cell lines supports the hypothesis of a preferential binding between L1 proteins and SINEs with high immunogenicity due to a higher dsRNA force. To test this hypothesis, **R**NA **I**mmuno**P**recipitation **Seq**uencing (RIP-Seq) assays for both α-ORF1p and α-ORF2p of L1 proteins were conducted in the embryonal carcinoma cell line N2102p (**Fig. 5E**), with detectable expression of ORF2p, a L1 ORF encodes the reverse transcriptase and endonuclease^41^. Transcripts enriched by co-IP were determined by differential expression analysis of RNA immunoprecipitated vs. total RNA and matched mock IP controls (**Methods**). L1 protein binding showed a specific preference to LINEs and SINEs: repeats that co-IPed with both ORF1p and ORF2p are mostly composed of LINEs and SINEs as expected (**Fig. S6C**). Overall, both SINE and LINE transcripts that bound ORF1p and ORF2p (Log2FC > 3, p.adj < 0.05) had significantly higher dsRNA force compared to the non-binding transcripts from the same repeat class, in contrast to DNA and LTRs (**Fig. 5F**). AluS comprises the majority of ORF1p and ORF2p binding SINEs (42%) (**Fig. 5G** & **Fig. 5H**). By comparing the composition of the SINEs that bound both ORF1p and ORF2p with those that did not bind, we found that AluYs, the youngest SINE family member, are exclusively bound by both L1 proteins (**Fig. 5G**).

### Tumors converge on L1 or ADAR1 to suppress immunogenic SINEs

Several studies have demonstrated that ADAR1, an RNA deaminase enzyme, can suppress immunogenic double-stranded RNAs through Adenosine-Inosine editing, with inverted SINEs being a predominant substrate^9, 26, 42–46^. To better understand the interplay between L1, ADAR1, and their impact on immunogenic SINEs in tumor cells, we examined ADAR1 activity from both transcriptional and functional perspectives. We found that samples with mutated *TP53* exhibit compromised expression of ADAR1, similar to the observed pattern for MDM2, a *TP53* suppressor gene (**Fig. 6A**). However, no significant differences in immunogenic SINE expression (AluS, AluY, SVA) were found under different *TP53* status (**Fig. 6B**). As expected, both ADAR1 expression and L1 insertion are anti-correlated with young/immunogenic SINEs (**Fig. S7A**). Interestingly, ADAR1 expression and L1 activity are inversely correlated with each other (**Fig. S7A**), suggesting that ADAR1 governs immunogenic SINE expression independently of L1.

**Figure 6.**
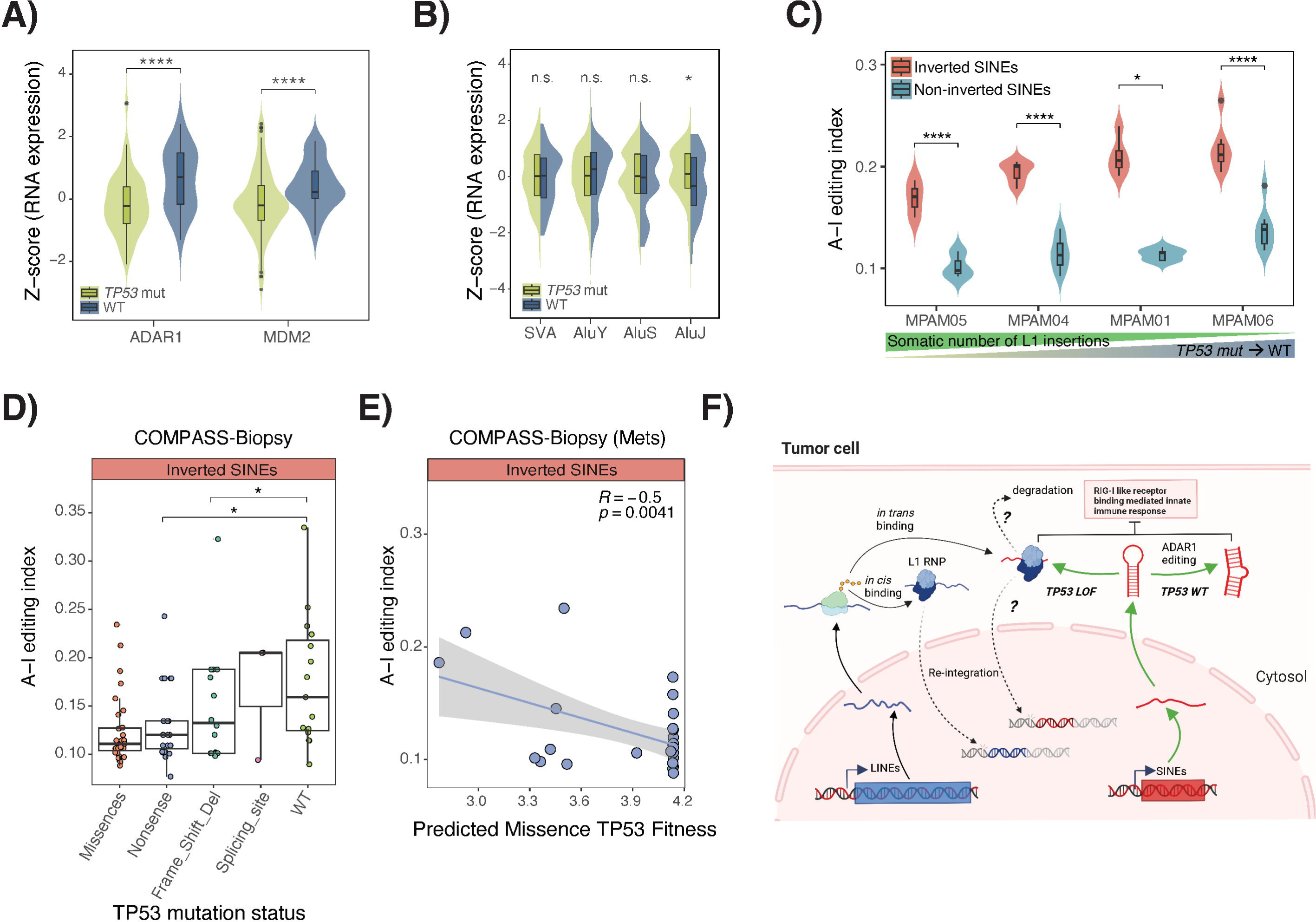
ADAR1 expression and function are dampened in *TP53* mutation. **A)** Violin plot shows the expression of ADAR1 and MDM2 in different *TP53* backgrounds in the autopsy PDAC cohort (p-values calculated by Wilcoxon test are labeled). **B)** Violin plot shows the expression of SINE subgroups in different *TP53* backgrounds in the autopsy PDAC cohort (p-values calculated by Wilcoxon test are labeled). **C)** A-I editing index for four patients with known different somatic mutation of *TP53* and number of somatic L1 insertions. Inverted SINEs are SINE copies with an adjacent copy next in the opposite direction that can potentially form dsRNA when expressed; non-inverted SINEs are SINE copies that do not meet these criteria (**Methods**). p-values out of Wilcoxon test are labeled. **D)** A-I editing index for PDAC biopsy samples in COMPASS trial stratified by different *TP53* mutation backgrounds. p-values calculated by Wilcoxon test are labeled. **E)** Scatter plot of A-I editing index as a function of predicted *TP53* missense mutation fitness^46^. **F)** Schematic diagram shows different strategies tumor cells leverage to maintain pro-tumorigenic SINE expression in *TP53* WT (via ADAR1 editing) and *TP53* mutant (via L1 binding) backgrounds. (* p < 0.05, ** p < 0.01, *** p< 0.001, **** p < 0.0001, representing the corresponding statistical significance)

To investigate the functional impact of ADAR1 on dsRNAs, we quantified A-G and T-C mismatched in RNAseq data as a readout of A-I editing. As predicted, inverted SINEs (**Methods**), have significantly higher A-I editing index compared to non-inverted SINEs in multiple patients (**Fig. 6C**). Interestingly, the A-I editing index is positively correlated with the total number of somatic L1 insertions and negatively correlated with *TP53* mutation status. For example, samples with more significant loss of function *TP53* mutations tend to have more somatic L1 insertions and lower A-I editing index (**Fig. 6C**). Stratifying samples based on different *TP53* mutation status revealed that patients with attenuated *TP53* mutations have higher A-I editing index in both primary and metastatic samples, with a stronger difference observed in metastasis (**Fig. S7B**). We confirmed this association in the independent cohort from the COMPASS trial^37^. As the COMPASS trail has a smaller number of reads mapped to SINEs due to its poly-A RNA-sequencing protocol (**Fig. S5B**), we see less A-I editing index compared to the PDAC autopsy cohort (**Fig. S7C**). However, the discrepancy between inverted and non-inverted SINEs was recapitulated (**Fig. S7C**). We then compared ADAR1 functionality in samples with different *TP53* mutation statues and found that samples with weaker *TP53* mutations or wildtype status had significantly higher A-I editing index than those with nonsense or frameshift deletions (**Fig. 6D**). Interestingly, primary samples had significantly higher A-I editing index compared to metastatic samples (**Fig. S7D**), which is opposite to the observed somatic L1 insertion trend (**Fig. S3D**). When comparing the A-I editing index of inverted SINEs with the number of somatic L1 insertions, we found a significant negative association in the rapid autopsy cohort (**Fig. S7E**). Those observations indicate that L1 mobility and ADAR1 functionality are complementary and mutually exclusive to each other.

To further explore the relation between ADAR1 functionality and *TP53* missense mutations more granular in the COMPASS trial, we used a previously developed method to compute the fitness of different *TP53* missense mutations based on their oncogenicity^47^. We observed a significant negative correlation between predicted missense mutation fitness (oncogenicity) and A-I editing index in metastatic samples, further demonstrating ADAR1 functionality is dependent on the oncogenicity of *TP53* mutations (**Fig. 6E**). Taken together, our findings suggest that L1 protein can repress immunogenic SINEs while, in parallel, ADAR1 can edit SINEs and ultimately lead to RNA degradation as an alternative immune surveillance mechanism when *TP53* is largely functional (**Fig. 6F**). These parallel mechanisms of immunogenic SINE regulation are essentially mutually exclusive and converge on maintaining immunogenic SINE expression in a pro-tumorigenic manner in PDAC.

## DISCUSSION

In this study, we show the interplay between two classes of retrotransposons, and how their interaction is influenced by *TP53* mutation status and RNA editing along tumor evolution in PDAC. Our study revealed a core **R**IG-**I L**ike **R**eceptor (RLR) associated IFN signature associated with the expression of immunogenic SINE elements, such as the AluY and AluS families (**Fig. 2A**). We computationally validated the mechanism whereby the transcripts of these younger Alu repeats have a higher likelihood to form dsRNA (**Fig. 2D**)^25^, acting as viral mimics that engage pattern recognition receptors, such as RIG-I and MDA5, which then upregulate immune responses^48^. We contend that one mechanism behind chronic inflammation is convergent evolution via multiple pathways on the regulation of immunogenic SINEs in cancer, and therefore the regulation of SINE expression could represent a multiple therapeutic targets^4^. To gain proliferative advantage a tumor cell has to balance a tradeoff between oncogenic potential and immunogenic cost^47^. Initially increased epigenetic plasticity in a tumor will trigger the acute expression of repeats which activate immune signaling^49^. SINE-induced innate immune responses will likely be initially detrimental to the tumor, and thus tumors adapt to keep SINE expression in check and shift to chronic pro-tumorigenic inflammation mediated by other pathways, such as DNA sensing pathways cGAS/STING^49^.

*De novo* L1 insertions in tumor samples were associated with the status of *TP53* mutations (**Fig. 3D**), supporting active differential surveillance of L1 mobility by mutant *TP53* in tumor evolution in addition to the repressive role of *TP53* on L1 activity^38, 51^. Our patient data support an antagonistic relationship between L1 activity and immunogenic SINE expression, in that high L1 insertion rates correlate with low immunogenic SINE expression (**Fig. 4B**). Consistently in PDAC cell lines, reducing the expression of L1 ORF1p by shRNA resulted in an increase of SINE expression. Our findings revealed a previously undescribed role for L1 in repressing anti-tumorigenic immunity elicited by SINEs. This hitherto unappreciated mechanism distinctly regulates immunogenic SINE expression in PDAC evolution: tumors evolve to escape the immunosurveillance by modulating the abundance of SINE RNAs via increased L1 activity. In contrast, for tumors with low L1 activity, ADAR1 keeps immunogenic SINEs in check via RNA editing. ADAR1 editing activity was highest in samples with wild type *TP53* and low L1 insertions and, conversely, lowest in samples with mutated *TP53* and high L1 insertions. We therefore propose a model whereby pancreatic tumors converge on regulation of dsRNA viral mimics by modulating their concentration via either L1-mediated removal or ADAR1 editing (**Fig. 6F**). Reduced levels of SINE-derived dsRNAs will result in reduced RLR mediated acute inflammation while shaping an immune suppressive environment potentially by facilitating M2 macrophage infiltration (**Fig. S4F**) that is likely a result of chronic rather than acute inflammation.

One of the major challenges in the treatment of PDAC is the highly immunosuppressive tumor microenvironment that is characterized by desmoplasia, which creates a mechanical barrier around the tumor, associated with chronic inflammation^52^. Chronic inflammation in cancer cells can promote resistance to immune checkpoint blockade (ICB), and this inhibition can be abrogated by remodulating IFN-I signaling in cancer cells^50^. Recent work proposed the combination of epigenetic therapy to induce viral mimicry, particularly the expression of immunogenic dsRNAs from inverted Alu elements, with the depletion of ADAR1, to prevent dsRNA editing^26^. Our study introduces the potential of inhibiting L1 activity to maintain immunogenic SINE expression. We thus envision a combination therapy that effectively targets PDAC tumors based on the status of their *TP53* mutations, L1 insertions and ADAR1 activity, with the goal to increase immunogenic SINE-derived dsRNA to trigger an effective immune response and avoid low level, chronic activation of inflammatory signaling. In summary, our results shed light on the critical evolutionary role of genomic “dark matter” – how repeats enable tumors to leverage viral-like adaptive responses to manipulate immunogenic vulnerabilities to their selective advantage.

## Methods

### Ethics statement

This study was approved by the Review Boards of Memorial Sloan Kettering Cancer Center and Massachusetts General Hospital.

### PDAC cell lines

PDAC3, PDAC6, PDAC8, PDAC9, were generated from metastatic ascites fluid of pancreatic adenocarcinoma patients at the Massachusetts General Hospital (MGH) under a discarded tissue protocol in accordance with the Massachusetts General Hospital IRB protocol 2011P001236 and Dana-Farber Harvard Cancer Center IRB protocol 02-240 as previously described^53^. PANC-1 (CRL-1469), MiaPaCa-2 (CRL-1420), Capan-1 (HTB-79), KP4, YAPC, DAN-G and BxPC-3 (CRL-1687) were purchased from the American Type Culture collection (ATCC).

### Western blot

PDAC cell lines were cultured into ultra-low attachment 6-well plate (Corning, USA, #3471) for 1 week. Cells were collected, washed, and lysed with lysis buffer containing 50 mM Tri-Cl (pH 6.8), 10% glycerol, 2% sodium dodecyl sulfate, 1 mM 1,4-dithiothreitol and Halt™ Protease and Phosphatase Inhibitor Cocktail (ThermoFisher Scientific, USA, #78440). The cell lysates were subjected to sodium dodecyl sulfate–polyacrylamide gel electrophoresis (SDS-PAGE) and then transferred to a 0.45 µm polyvinylidene difluoride (PVDF) membrane (Millipore, MA, USA, #IPVH00010). The membranes were blocked with 3% of bovine serum albumin (BSA, Sigma-Aldrich, # A2058) for 1 hour and applied with primary antibodies against LINE1 ORF1p (Millipore, USA, #MABC1152) and β-actin (Cell Signaling Technology, USA, #4970S) for overnight at 4^◦^C. Membranes were washed with 1X phosphate-buffered saline (PBS) containing 0.1% tween-20 (Sigma-Aldrich, USA, #P1379) (PBST) and incubated with horseradish peroxidase (HRP)-conjugated secondary antibodies (HRP-conjugated goat anti-rabbit (Cell Signaling Technology, # 7074S) and HRP-conjugated goat anti-mouse (Cell Signaling Technology, # 7076S). Signal was detected with enhanced chemiluminescence (SuperSignal™ West Pico PLUS Chemiluminescent Substrate, Thermo Scientific, #34577), and images were developed using G:BOX (SYNGENE).

### Total RNA preparation and sequencing

#### PDAC autopsy cohort

Raw bam files of RNAseq data were kindly provided by the Iacobuzio lab associated with their study^18^. Briefly, frozen sections were cut from samples for histological review and regions of interest were microdissected for extracting total RNA using TRIzol (Life Technologies) followed by Rneasy Plus Mini Kit (Qiagen). Each RNA sample was initially quantified by Qubit 2.0 Fluorometer (Thermo Fisher Scientific). Samples were additionally quantified by RiboGreen and assessed for quality control using an Agilent BioAnalyzer in the Integrated Genomics Core at MSKCC and 513 ng to 1.0 µg of total RNA with an RNA integrity number ranging 1.3–8.3 underwent ribosomal depletion and library preparation using the TruSeq Stranded Total RNA LT kit (Illumina, RS-122-1202) according to instructions provided by the manufacturer with eight cycles of PCR. Samples were barcoded and run on a HiSeq 4000 in a 100 bp per 100 bp or 125 bp per 125 bp paired end run, using the HiSeq 3000/4000 SBS kit (Illumina). On average, 94 million paired reads were generated per sample and 26% of the data were mapped to the transcriptome.

#### PDAC cell lines

Cells were cultured in 3D tumoursphere condition for 7 days then, cells were lysed, and total RNA was extracted using the miRNeasy Mini Kit (QIAGEN, cat# 217004). Libraries were prepared for sequencing using the SMARTer Stranded Total RNA-seq Kit v2 (cat# 634413) – Pico Input Mammalian (Takara Bio USA). RNA samples were first synthesized into cDNA, and adaptors for Illumina sequencing were added using PCR. Then, the PCR products are purified using AMPure Beads, and ribosomal cDNA was removed. After depletion, the samples were further amplified using PCR, then purified again using AMPure Beads. Each sample was then qPCR quantified using KAPA Library Quantification Kit (KAPA Biosystems). Samples were then pooled together and quantified again using the KAPA Library Quantification Kit. The Pooled library was then sequenced using the NextSeq 1000 Sequencing Kit.

### Mapping

To re-align sequences to our pipeline, we first converted all bam files into fastq using *samtools v*1.9. RNA sequencing reads were 3’ trimmed for base quality less than 20 using *skewer*. Reads shorter than 40 after trimming were removed. Then the quality score of the remaining bases were sorted, and the quality at the 20th percentile was computed. Reads with quality at the 20th percentile less than 15 were discarded. Only paired reads which both pass the quality check were mapped to the reference genome (hg38) using *STAR v*2.7 with default parameters. Gene counts were assigned based on Gencode annotation using *featureCounts* (*subread* − 1.4) with the external Ensembl annotation. Repeats counts per element family were primarily quantified against RepeatMaster using *featureCounts* (*subread* − 1.4) and then adding the counts of the unassigned reads that mapped to Repbase consensus sequence. Repeat counts of a given family is the sum of mapped reads to RepeatMassker and unmapped reads against Repbase.

### Counts filtering and normalization

Expression of repeats and coding genes was normalized across samples using trimmed-mean of M-values (TMM) in *edgeR*. The size factor for each sample was calculated using *calcNormFactors* based on coding genes alone as described in the previous study^19^. Low-count gene/repeats in each sample with counts per million (CPM) smaller than 2 were removed before calculating the size factor. Log2 transformed CPM were used for downstream visualization and correlation analysis.

### RNA based tumor purity estimation and differential expression analysis

*ESTIMATE v*2.0.0^20^ in R was used to infer the tumor purity from total transcriptome data in 214 human samples. Differential expressions of coding genes and repetitive elements were analyzed separately using *DESeq*2 *v*1.33.4^54^ by adding tumor purity as a covariate in formula design to minimize the bias caused by high stromal contamination in human samples or by default parameter for PDAC or RPE cell lines.

### Gene set enrichment analysis (GSEA)

GSEA was applied on a gene list ranked by the differential expression on the basis of the methods described^55^. To test the gene set significance, an enrichment score is defined as the maximum distance from the middle of the ranked list. Gene set with positive enrichment score indicates upregulation with a positive log2FC, whereas the negative enrichment score is suggesting a downregulation with a negative log2FC. *clusterProfiler v*4.0^21^ was used to run this analysis against HALLMARK pathways in MSigDB collection.

### dsRNA force and evolutionary distance calculation for each SINE copy in hg38

By searching for a pairable SINE (nearby SINE in the opposite direction) located upstream or downstream of the target SINE within a 3000-bp window. We determined the likelihood of discovering a suitable matching copy and corresponding matched length between two SINEs

### dsRNA force calculation

Given a set of nucleotide frequencies *f*(*σ*), the probability of observing a pair of complementary nucleotides is

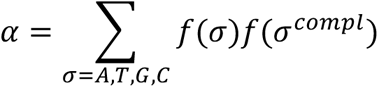

From this quantity it can be computed the expected length *N_ds_*(*L*, 0) of the longest fully complementary strand of a sequence composed by L nucleotides sampled from the frequency distribution *f*(*σ*), obtaining after some approximations^25^

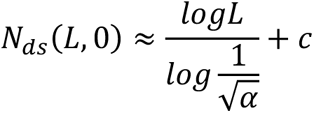

where for the correction constant we obtain c = −2.2 when considering both canonical Watson-Crick pairs and Wobble pairs as complementary basepairs. For a given RNA sequence, the dsRNA force *x*_*ds*_ is introduced to fix mismatches from the expected values *N_ds_*(*L*, 0) and the observed ones, 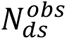 Following Sulc et al., we obtain the equation

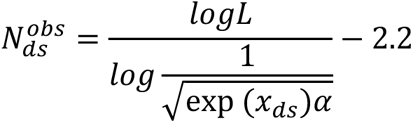

that can be used to compute the dsRNA force *x_ds_*. for an RNA sequence of length L having length of the longest complementary strand equal to 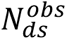.

In the case of SINE element, we are interested in computing the dsRNA force when one of the two complementary segments composing the longest complementary strand is constrained to be within the SINE insertion. To do that, we consider windows of 3 kb around the insertion, and use the equation

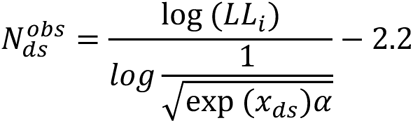

where *L_i_* is the insert length. This equation is obtained again by following the approach used by Sulc et al.^25^, but considering that one of the two strands of the longest double-strand segment must be in a subsequence of length *L_i_*.

### RNA immunoprecipitation sequencing and data analysis

#### RNA extractions and coimmunoprecipitations (RIP) from EC cells – N2102Ep

Embryonal carcinoma cells were cultured at 37 °C in humidified incubators maintained with 7% CO2 atmosphere. N2102Ep Clone 2/A6 cells (Merk, #06011803) were cultured in DMEM (high glucose, no sodium pyruvate; Thermo Fisher, #11965092), supplemented with 10% (v/v) FBS, 1xpenicillin/streptavidin and 2 mM Glutamine (Thermo Fisher, #25030024). Largescale growth, harvesting, cryomilling, and coIP was achieved as previously described^56–58^, summarized as follows. αORF1p and αORF2p targeted coIPs used inhouse made^59^ magnetic affinity media: for αORF1p [15 μg antibody / mg magnetic beads], we used the 4H1 monoclonal antibody (Millipore Sigma, #MABC1152); for αORF2p [10 μg / mg magnetic beads] we used the clone 9 monoclonal antibody^41^. CoIPs were conducted using 100 mg cell powder, extracted at 25% (w:v) in 20 mM HEPES pH 7.4, 500 mM or 300 mM NaCl, 1% (v/v) Triton X100, 1x protease inhibitors (Roche, #1187358001), and 1:250 RNasin (Promega, #2515). Centrifugally clarified cell extracts were incubated with affinity medium (20 μl of slurry for αORF1p and αORF2p for 30 min at 4 °C. The solutions were made with nucleasefree H_2_O and experiments were conducted using nucleasefree tubes and pipette tips. Macromolecule extractions performed on these cell lines as described typically yielded between 450∼500 μl of soluble extract at 6∼8 mg/ml of protein as assessed by Bradford assay (Thermo Fisher, #23200). After target capture, washing the media was performed with the same solution without protease inhibitors and with RNase at 1:1000. RNAs were eluted from the affinity media after RIP with 250 μl of TRIzol Reagent (Thermo Fisher, #15596026). After adding chloroform to the TRIzol eluate, the separated aqueous phase (containing RNAs) was obtained using Phasemaker tubes (per manufacturer’s instructions; Thermo Fisher, #A33248), and was then combined with an equal volume of ethanol and further purified using a spin column according to the manufacturer’s instructions (Zymo Research, #R2060). Eluates from αORF1p and αORF2p coIPs were not treated with DNase I on-column, this was done during the sequencing library preparation (described, below). Purified nucleic acids from αORF1p and αORF2p coIPs were eluted in 6ul of nucleasefree water; in all cases 1 μl was used for quality analysis and the remainder conserved for RNAseq. Mock RNA coIP controls were prepared in an identical manner using naïve polyclonal mouse IgG (control for αORF1p; Millipore Sigma, #I5381) or naïve polyclonal rabbit IgG (control for αORF2p: Innovative Research, #IRBIGGAP10MG). Total RNA controls were prepared by combining up to 35 μl of the clarified cell extracts with up to 500 μl of Trizol, vortex mixing for 1 min, then snap freezing in liquid nitrogen and then later proceeding as above.

#### cDNA library preparation and RNA-seq

All the sequenced samples/replicates that are reported in this study are listed in **Table S4**. αORF1p and αORF2p RIPseq: RNA extractions were quantified, and quality controlled using RNA Pico Chips (Cat. #50671513) on an Agilent 2100 BioAnalyzer. RNA-seq cDNA libraries were prepared 15 using the Trio RNA-seq Library Prep kit (Tecan, #0357A01) with Any Deplete Probe MixHuman rRNA (Tecan, #S02305). DNase treatment preceded cDNA synthesis. cDNA synthesis: 3 5ng of input RNA from αORF1p RIPs and mock IPs, 1 ng from αORF2p RIPs and mock IPs, and 50 ng of total RNA were used with 8 (2+6) cycles of pre-depletion PCR library amplification and 8 (2+6) cycles of post-depletion amplification; the libraries were purified using Agencourt AMPure XP beads (Beckmann Coulter), quantified by qPCR, and the size distribution was checked using the Agilent TapeStation 2200 system. Final libraries were sequenced, pair-end, at 50 bp read length on an Illumina NovaSeq 6000 v1.5 with 2% PhiX spike-in.

#### RIPseq read mapping and quantification

Reads were trimmed and quality checked using skewer^60^. Briefly, ends of the reads were trimmed to remove Ns and bases with quality less than 20. After that, the quality scores of the remaining bases were sorted, and the quality at the 20th percentile was computed. Reads were discarded if its quality at the 20th percentile was less than 15. In addition, reads shorter than 40 bases after trimming were discarded. If at least 1 of the reads in the pair failed the quality check and had to be discarded, we discarded the mate as well. Quality filtered reads were mapped to annotated repeat loci in RepeatMasker using software: Quantifying Interspersed Repeat Expression (SQuIRE) (https://github.com/wyang17/SQuIRE)61. The SQuIRE pipeline first obtains reference annotation files from RepeatMasker, then aligns reads using STAT, and lastly, quantify locus specific repeat expression by redistributing multimapping read fractions in proportion to estimated TE expression with an expectation maximization algorithm.

#### Selection of RIPSeq enriched repeats/transcripts

Targeted protein enriched transcripts were selected by Log_2_(CoIP/mock) > 3, Log_2_(CoIP/ totalRNA) > 3, and Benjamini-Hochberg adjusted p-value < 0.05. Similarly target protein depleted transcripts were selected by Log_2_(CoIP/mock) < 3, Log_2_(CoIP/total RNA) < 3, and Benjamini-Hochberg adjusted p-value < 0.05.

### DNA sequencing and mutation calls

Genomic DNA was extracted from each tissue using QIAamp DNA Mini Kits (Qiagen). WGS, WES and alignment were performed as previously described^62^. Briefly, an Illumina HiSeq 2000, HiSeq 2500, HiSeq 4000 or NovaSeq 6000 platform was used to target a coverage of 60× for WGS samples and 150× for WES samples. The resulting sequencing reads were analyzed in silico to assess quality, coverage, as well as alignment to the human reference genome (hg19 or hg38) using BWA^63^. All WES samples and WGS in MPAM samples were aligned to hg19, WGS data from PAM samples were aligned to hg38. After read deduplication, base quality recalibration and multiple sequence realignment were completed with the PICARD Suite and GATK v.3.1 (refs. 52,53) somatic single-nucleotide variants and insertion–deletion mutations were detected using Mutect v.1.1.6 and HaplotypeCaller v.2.4^64, 65^. We excluded low-quality or poorly aligned reads from phylogenetic analysis. Filtering of called somatic mutations required each mutant to be observed in at least one neoplastic sample per patient with at least 5% variant allele frequency and with at least 20× coverage; correspondingly, each mutant had to have been observed in <2% of the reads (or fewer than two reads in total) of the matched normal sample with at least 10× coverage.

### Evolutionary analysis

We derived phylogenies for each set of samples by using Treeomics 1.7.9^39^. Each phylogeny was rooted at the matched patient’s normal sample and the leaves represented tumor samples. Treeomics employs a Bayesian inference model to account for error-prone sequencing and varying neoplastic cell content to calculate the probability that a specific variant is present or absent. The global optimal tree is based on mixed integer linear programming. All evolutionary analyses were performed on the basis of WGS depending on the sequencing data availability (**Figure 1A**)^62^. Somatic alterations present in all analyzed samples of a PDAC were considered clonal, in a subset of samples or a single sample considered subclonal.

### Whole Genome Inverted Repeats Analysis

We applied the inverted-repeat pipeline^4^ on all repeat elements in the human hg38 genome by considering all repeat elements in the RepeatMasker list of elements which was downloaded from the UCSC database. All repeat loci that are within 3Kb distance were paired and aligned locally in the reverse complement manner by the Smith–Waterman algorithm using the exonerate tool (v.2.2.0)^66^. Two different penalty scores were applied for gap opening and extension which are a 12-penalty score for gap opening and a 4-penalty score for gap extension. Aligned pairs that are separated by distance greater than 3Kb or pairs that are aligned with alignment score less than 500 were discarded. There are about 795,058 repeat pairs that were aligned as inverted repeats pairs. Among these pairs that were aligned as inverted repeats, there are about 750,004 pairs formed by Alu elements.

### Calling L1 retrotransposition

We called L1 retrotranspositions using TotalREcall^35^ with matched normal samples to retain somatic mutations. Overlapping events found in normal samples, were removed. To confirm the reliability of calls and remove remaining false-positive events we visually inspected all called *de nova* L1 insertion candidates focusing on two supporting pieces of evidence: (1) poly-A tails and (2) target site duplications/deletions using Integrative Genomics Viewer. Additionally, we excluded variants with a low number of supporting reads (fewer than 4 reads) to exclude potential artefacts.

### A-I editing index

All possible RNA editing events were called using REDItools2.0^67^ against all annotated SINE loci in hg38. The estimated RNA editing events were selected for single- and multi-A–I nucleotide substitutions. All subset RNA editing profiles were then subjected to the following quality control measures to remove editing estimates that fell below the following thresholds: minimum average quality score (MeanQ) of 25; minimum depth of coverage of 10; and minimum average editing frequency of 1%. Only non-mitochondrial edits were subject to downstream analysis for clarity. The resulting subset of A–I editing events was used to calculate an A-I RNA Editing Index using the *RNAEditingAnalysisTools* R package^68^. Here, a normalized sample-specific global A-I RNA Editing index was calculated as the total proportion of edited reads over reads mapping to edited sites (Editing Reads/Coverage).

### Histology, IHC and IF

#### LINE-1 ORF1p Immunohistochemistry

Tissues were fixed overnight in 10% formalin, embedded in paraffin, and cut into 5 µm sections. Slides were heated for 20 min at 55°C, deparaffinized, rehydrated with an alcohol series, and subjected to antigen retrieval with citrate buffer (Vector Laboratories Unmasking Solution, H-3300) for 15 min in a pressure cooker set on high. Sections were treated with 3% H_2_O_2_ (in PBS) for 15 min followed by a wash in deionized water, washed in PBS, then blocked in 5% BSA (in PBS). The following primary antibody was incubated overnight at 4°C in blocking buffer: LINE-1 ORF1p (ab230966, Abcam, 1:200). Vector ImmPress anti-Rabbit HRP and ImmPact DAB kits (Vector Laboratories) were used for secondary detection. Tissues were then counterstained with Haematoxylin, dehydrated and mounted with Permount (Fisher). Images were acquired on a Axio Imager Z2 platform (Zeiss).

#### LINE-1 ORF1p and dsRNA Immunofluorescence

PDAC cell lines were plated onto the Collagen type-I, rat-tail (50 µg/ml) coated 12 mm coverslips and allowed to adhere for at least 48 hrs before use in experiments. Cells were fixed for 15 min with freshly prepared 4% (w/v) paraformaldehyde in PBS, permeabilized with 0.5% (v/v) Triton X-100 in PBS for 10 min. Then, cells were blocked with 2% BSA solution for 1 hr, incubated with primary antibodies (anti-dsRNA, clone rJ2; Sigma-Aldrich, cat# MABE1134 and anti-LINE-1 ORF1p, clone 4H1; Sigma-Aldrich, cat# MABC1152) with desired concentration (1:50) in blocking solution for 1 hr at room temperature. After three washes with 0.1% Triton X-100 in PBS (PBST) were further incubated with fluorophore-conjugated secondary antibodies (goat anti-Mouse IgG2a Cross-Adsorbed Secondary Antibody, Alexa Fluor 594, Thermo Scientific, cat# A-21135, goat anti-Mouse IgG1 Cross-Adsorbed Secondary Antibody, Alexa Fluor 488, Thermo Scientific, cat# A-21121) for 1 hr at room temperature. Following additional three washes with PBST, DAPI solution was applied for 1 min then washed. Coverslips were mounted in Fluoromount-G® Mounting Medium (SouthernBiotech, cat# 0100-01) and imaged by fluorescence microscopy (Nikon). The fluorescence intensity of dsRNA and LINE-1 ORF1 was measured using Image J software.

### LINE1 ORF1p shRNA knocking down

#### Plasmid construction and generation of stable cell line

shRNA sequences targeting LINE-1 ORF1^69^ were cloned into pLKO.1-TRC cloning vector (Addgene, cat# 10878) using EcoR1 and AgeI restriction enzyme digestion. For lentiviral production, HEK293T cells were seeded at 80% density in a 10-cm tissue cell culture treated dish and transfected with the 6 µg of expression plasmid and packaging plasmids 2 µg of pMD2.G (Addgene, cat# 12259) and 4 µg of psPAX2 (Addgene, cat# 12260) using Lipofectamine 2000 Transfection Reagent according to manufacturer’s instructions. Conditioned medium containing recombinant lentivirus was collected and filtered through 0.45 µm filters. The LINE-1 ORF1 shRNA containing lentivirus medium was added to cells with 8 µg ml-1 of polybrene (Santa Cruz Biotechnology, cat# sc-134220). Approximately 48 hrs after infection, the cells were selected with 1 µg ml-1 puromycin (InvivoGen, cat# ant-pr-1).

#### Immunoblotting

PDAC cells were harvested, washed, and lysed with lysis buffer (50 mM Tri-Cl (pH 6.8), 10% glycerol, 2% sodium dodecyl sulfate, 1 mM 1,4-dithiothreitol and Halt™ Protease and Phosphatase Inhibitor Cocktail (ThermoFisher Scientific, USA, cat# 78440). Total protein concentration was determined by BCA protein assay (Thermo Fisher Scientific, cat# 23227). The cell lysates were performed to sodium dodecyl sulfate–polyacrylamide gel electrophoresis (SDS-PAGE) and then transferred to a polyvinylidene difluoride (PVDF, 0.45 µm) membrane (MilliporeSigma, MA, USA, cat# IPVH00010). The membranes were blocked with 3% of bovine serum albumin (BSA, Sigma-Aldrich, cat# A2058) for 1 hr and applied with primary antibodies for overnight at 4◦C. Membranes were washed with 1X phosphate-buffered saline (PBS) containing 0.1% Tween-20 (Sigma-Aldrich, cat# P1379) (PBST) for 10 minutes 3 times and incubated with horseradish peroxidase (HRP)-conjugated secondary antibodies. Signals were developed with enhanced chemiluminescence (SuperSignal™ West Pico PLUS Chemiluminescent Substrate, Thermo Scientific, cat# 34577), and images were detected using G:BOX (SYNGENE).

The following primary antibodies were used: anti-LINE-1 ORF1p (MilliporeSigma, cat# MABC1152), GAPDH (Cell Signaling Technology, cat# 2118) were used following the manufacturer’s suggested protocols.

### Statistics and reproducibility

All statistics and graphs were performed and made using *R v*4.0.3 and organized in adobe illustrator 2022. Parametric distributions were compared by a two-sided t-test, with correction using Fisher’s exact test for sample sizes <5. Nonparametric distributions were compared using Wilcoxon test and for analysis of contingency tables, a two-sided Fisher’s exact test was used. Each analysis is described in the Results. Statistical significance was considered if the adjusted P value (after Benjamini - Hochberg correction) was <0.05. The FDR q value was used for GSEA. No statistical method was used to predetermine sample size. No data were excluded from the analyses as long as the library and/or sequencing quality passed our criteria. The experiments were not randomized. The investigators were not blinded to allocation during experiments and outcome assessment except for review of histological slides.

## Data and code availability

Data availability RNA and DNA sequence data for this study have been deposited at the European Genome-phenome Archive under accession number EGAS00001003974. Original data will be made public upon acceptance. Code will likewise be deposited on GitHub.

All other data supporting the findings of this study are available from the corresponding author upon reasonable request.

## Acknowledgements

This research was funded in part through the NIH/NCI Cancer Center Support Grant P30 CA008748 (S.S., A.S., B.G.); NIH grants R01AI081848 (N.V., B.G.), R01CA240924 (A.S., B,G); Fondation de la Recherche Médicale: and U01CA228963 (S.S., A.S., B.G.); the V Foundation for Cancer Research (A.S.); and the Pershing Square Sohn Prize-Mark Foundation Fellowship (A.S., O.A.-W., N.V., B.G.). Canadian Institute of Health Research (CIHR), New Investigator salary award (201512MSH360794-228629 to D.D.C.). K.M.T. was supported by the Jane Coffin Childs Memorial Fund for Medical Research and a Shulamit Katzman Endowed Postdoctoral Research Fellowship. This work was supported by MSKCC’s David Rubenstein Center for Pancreatic Research Pilot Project (to S.W.L.) and NIH grant P01CA013106 (to S.W.L.). S.W.L. is an Investigator of the Howard Hughes Medical Institute and the Geoffrey Beene Chair for Cancer Biology at MSKCC. The authors would like to acknowledge Dr. Faiyaz Notta for sharing the COMPASS raw data, and the productive conversations with Dr. Wilson McKerrow, the Greenbaum laboratories, Ting laboratories and Burns laboratories and thank Nicole Rusk for reading and editing the manuscript.

## Author contributions

S.S., A.S., and B.G. acquired the data, conceptualized the work, designed experiments, visualized data, and interpreted results. A.S. designed all pipelines for data analysis. S.S. performed data analysis. S.S. wrote the manuscript and B.G. edited the manuscript. J.H. & C.A.I.-D. collected patient autopsy samples and performed the treeomics analysis and mutation calls from WGS data. E.Y. and D.T. designed and performed L1 KD, RNAseq sequencing, and IF in PDAC cell lines. K.M.T. & S.L. performed IHC in human samples. J.CB. performed ADAR1 editing index analysis. A.DG. performed genome wide dsRNA force, kimura distance calculation. D.H performed predicted *TP53* fitness. H.L. conducted the pipeline running and data management. H.J., H.L. and J.LC. generated RIPseq data from N21201p cell line. S.M. provided the annotation of inverted SINEs. N.V. & D.T. provide guidance in data analysis and experimental design. R.M., P.C., and A.K. collected recuts from FFPE samples from LWP.

## Extended Figure Legends

**Figure S1. A)** Scatter plot of tumor purity estimated based on RNAseq data (x-axis, using ESTIMATE) and DNAseq data (y-axis, using FACETS). **B) G**ene **S**et **E**nrichment **A**nalysis (GSEA) of HALLMARK gene sets. GSEA was applied to the gene list ranked by Log_2_FoldChanges between metastasis (n=90) and primary (n=124). The Log_2_FoldChanges have been corrected for tumor purity (**Methods**). **C)** Distribution of Pearson correlation coefficients between median expression of genes in each cluster and subfamily-based expression of repeats grouped and colored by classes. **D)** Scatter plot of correlation with RLR signature score as a function of base mean expression of different repeat classes. Spearman correlation coefficient and associated p value are labeled.

**Figure S2. A)** Heatmap of normalized and column-wise scaled SINE expression and RLR signature genes (C4) expression across all 214 samples. Color annotation for each sample (by row) is labeled for L1HS expression, tissue categories, patient ID, and sample site (primary/metastasis). 214 samples are clustered into three groups with either low (bottom cluster), medium (middle cluster), and high (top cluster) expression in L1HS, SINEs and RLR signature genes. **B)** Fraction of repeats from high (dsRNA force > 0.5) or low dsRNA force (dsRNA force < 0.05) regions in each evolutionary group. **C)** Violin plot comparing the correlation with RLR signature between SINEs with high and low dsRNA force (cutoff = 0.5). **D)** Scatter plot of immunogenicity of SINE elements (averaged correlation with RLR signature) and mean foldability (dsRNA force) of each SINE element across all loci in the genome^25^.

**Figure S3. A)** Immunohistochemistry staining for L1 ORF1p in samples from rapid autopsy cohort. The manually quantified IHC score are labeled on the example images. DAPI are used for nuclear staining. **B)** Scatter plot of manually quantified IHC score of ORF1p expression as a function of the number of somatic L1 insertions. **C)** Boxplot of number of L1 insertion in samples collected from primary and metastatic sites in PDAC biopsy cohort in COMPASS trail^36^. p-values of t-test significance are labeled. **D-E)** Boxplot of number of L1 insertion in samples with wild type and different types of *TP53* mutants in PDAC biopsy cohort in COMPASS trail. p-values after t-test are labeled. **F)** Integrative analysis of L1 insertion phenotypes and tumor evolution of patient MPAM05. Samples are boxed with colors representing the tissue type. Target site features is coded as red (target site duplication) and green (target site deletion).

**Figure S4. A)** Volcano plot of Log_2_FC of repeats between samples with high L1 insertions and low L1 insertions including three extreme samples from patient MPAM5. **B)** Volcano plot of Log_2_FC of RLR signature genes, defined in Fig. 1C are repressed in samples with high L1 activities. **C)** Spearman correlation coefficients between number of somatic L1 insertions and repeat expression grouped by repeat classes in autopsy PDAC cohort. **D)** Violin plot of spearman correlation between L1 insertions and expression of SINEs. SINE elements are colored based on their evolutionary time. **E)** Distribution of Pearson correlation coefficient between myeloid cells infiltration estimated using CIBERSORTx and repeats expression in different classes across 214 samples. **F)** Violin plot of Pearson correlation coefficient between absolute M2 Macrophage infiltration estimated using CIBERSORTx and different repeats expression. Benjamini-Hochberg corrected p-values of t-test are labeled. Pearson correlation between absolute M2 Macrophage infiltration and expression of different SINE subtypes. SINE subtypes are colored based on their evolutionary age: SINE2/SINE3, SINE1 (Monomeric and dimeric AluJ, AluS and AluY, and SVA).

**Figure S5. A)** Spearman correlation between L1 insertions and Myeloid cells infiltration. **B)** Barplot of ratios of reads mapped to SINEs and coding genes in COMPASS (n=203, polyA-selection) and autopsy PDAC cohort (n=214, rRNA depletion). **C)** Volcano plot of Log_2_FC of repeats with high L1 ORF1p and low L1 ORF1p expressing PDAC cell lines. **D)** Scatter plot of tumor purity corrected Log_2_FoldChange of all SINE elements in PDAC patients with different L1 insertions and their Log_2_FoldChange in PDAC cell lines with different L1 ORF1p expression.

**Figure S6. A)** Pearson correlation coefficients between L1 ORF1p expression normalized by GAPDH and repeat expression grouped by classes in L1 knocking down in cell line PDAC6. **B)** Volcano plot of Pearson correlation coefficient between SINE expression and ORF1p expression in PDAC6 cell line with different L1 knocking downs. **C)** Heatmap shows the normalized and scaled expression (Z-score) of L1 ORF1p and ORF2p binding repeats (colored by classes on columns) in the RIPseq experiment.

**Figure S7. A)** Heatmap of pairwise correlation among young SINE repeats (AluY and SVAs), L1 insertion, and expression of genes ADAR1 and ZBP1. **B)** A-I editing index of inverted SINEs for patients with known different somatic mutation of *TP53* stratified by tumor sites. **C)** A-I editing index for PDAC autopsy samples and COMPASS biopsy samples stratified by inverted and non-inverted SINEs. **D)** A-I editing index for PDAC biopsy samples in COMPASS trail stratified by tumor sites. **E)** Scatter plot of A-I editing index on inverted SINEs as a function of log10 based number of somatic L1 insertions. (* p < 0.05, ** p < 0.01, *** p< 0.001, **** p < 0.0001, representing the corresponding statistical significance of Wilcoxon test)

## Extended Table Legends

**Table S1.** List of 140 IFN and ISG genes with assigned clusters identified using co-expression hierarchical clustering.

**Table S2.** List of L1 insertions loci, supporting reads, and clipping reads information in PDAC autopsy samples

**Table S3.** Western blot and immunofluorescent quantifications of L1 ORF1p and dsRNA in PDAC cell lines.

**Table S4.** List of annotated samples subject for L1 ORF1p/ORF2p RIP-seq analysis. Salt concentration, cell line information, antibodies, replicates and sequencing detailed are listed.

