## Supplementary figures and images for "Cancer cells co-evolve with retrotransposons to mitigate viral mimicry"

### Figure S1.png

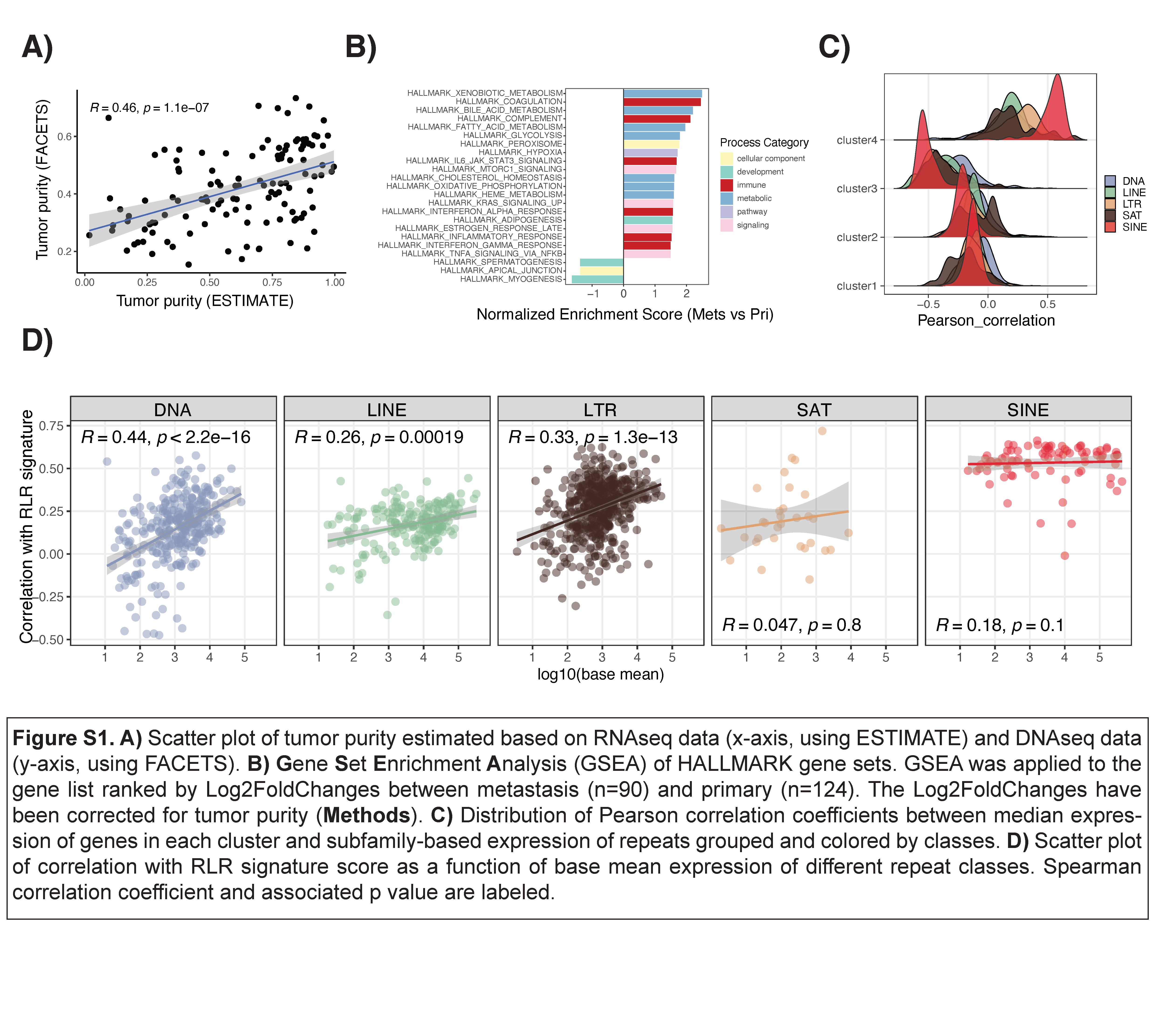

### Figure S2.png

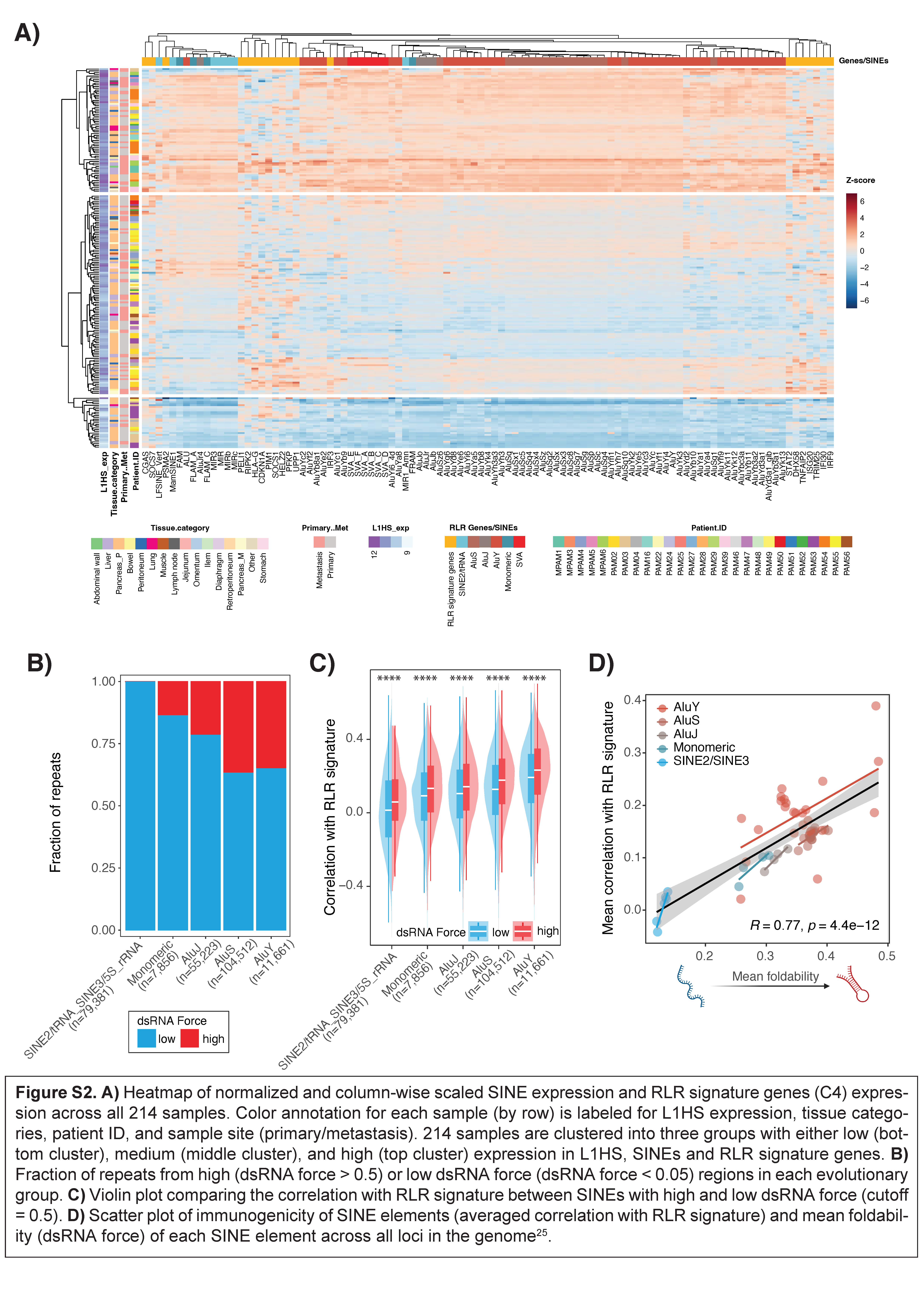

### Figure S3.png

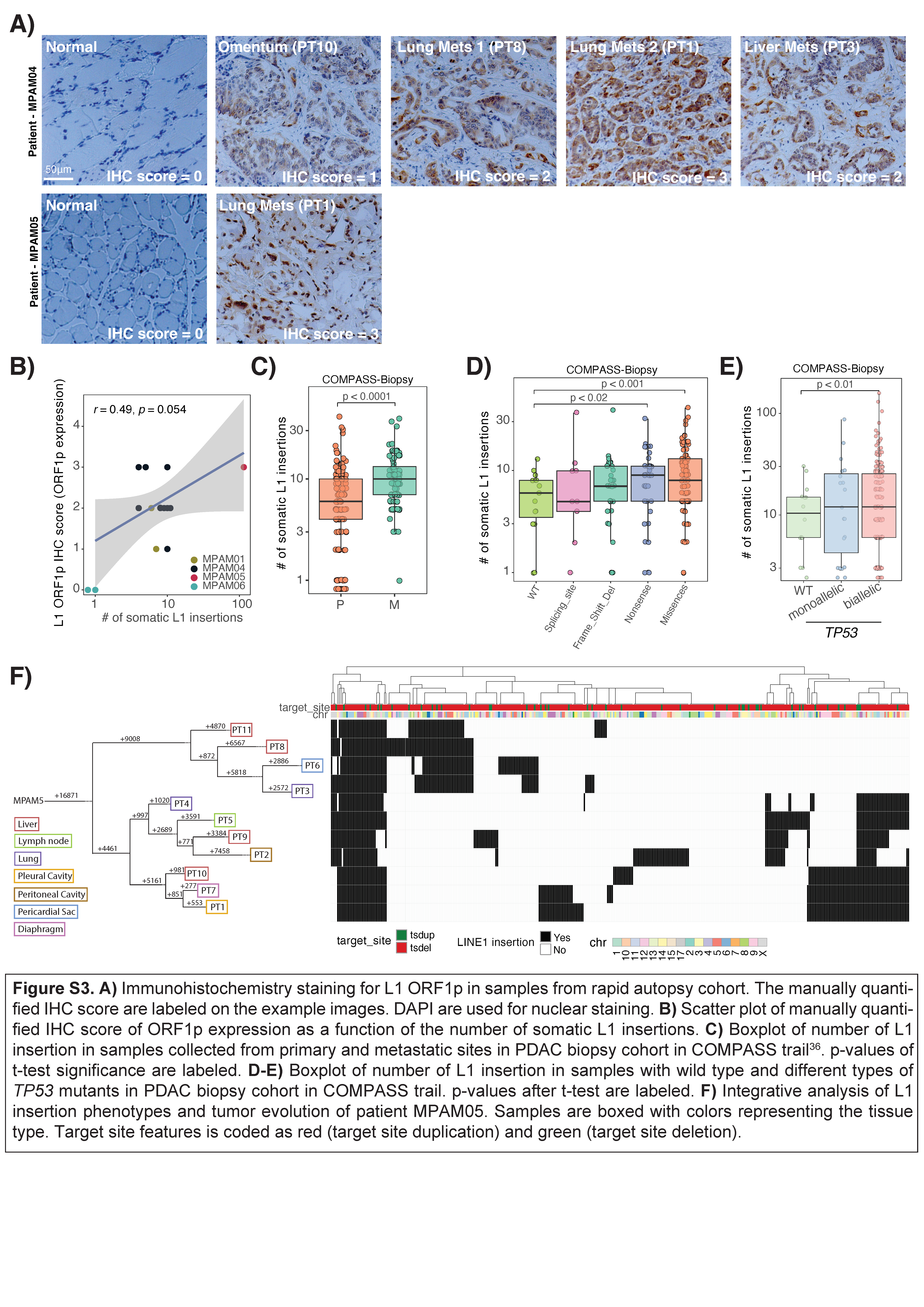

### Figure S4.png

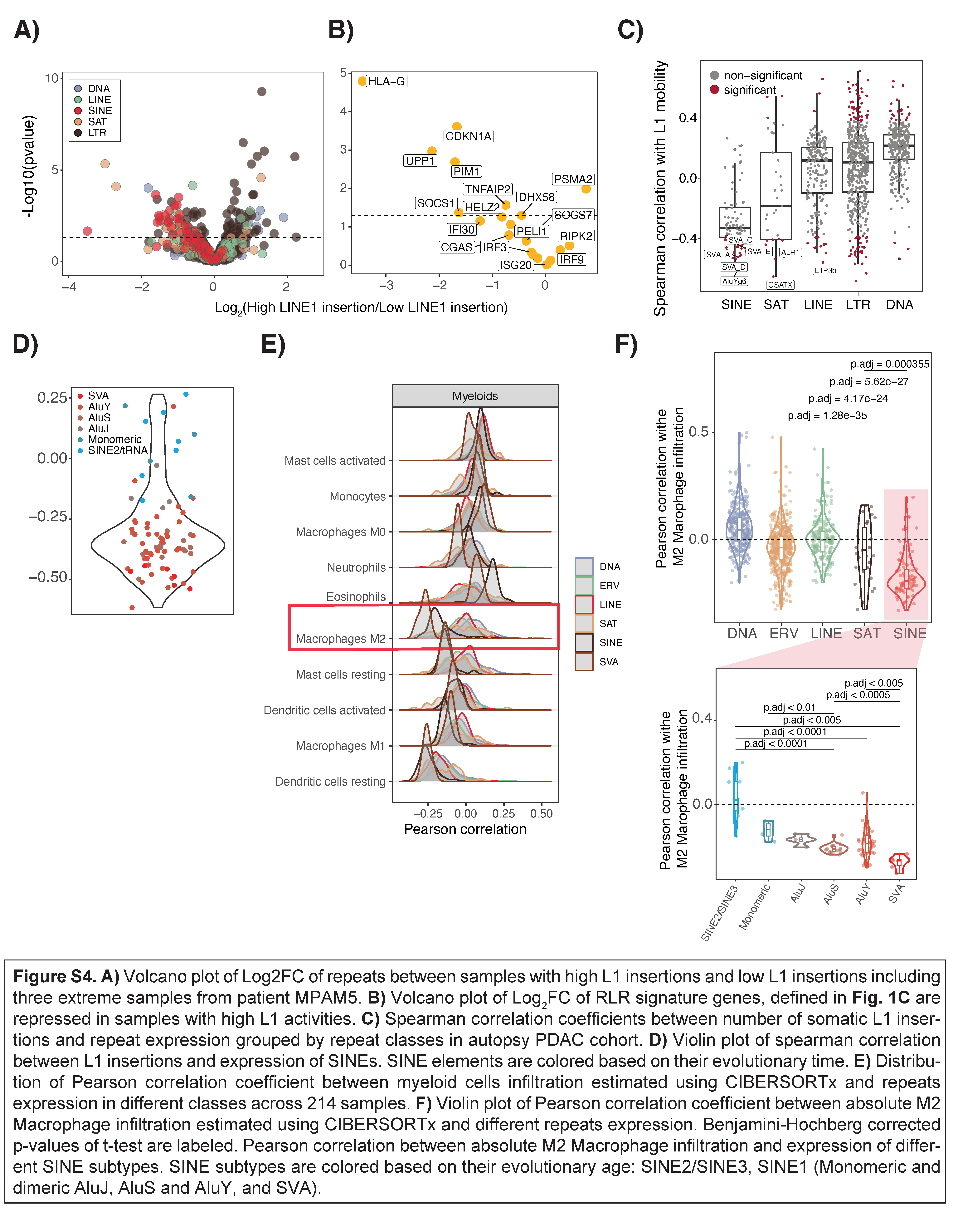

### Figure S5.png

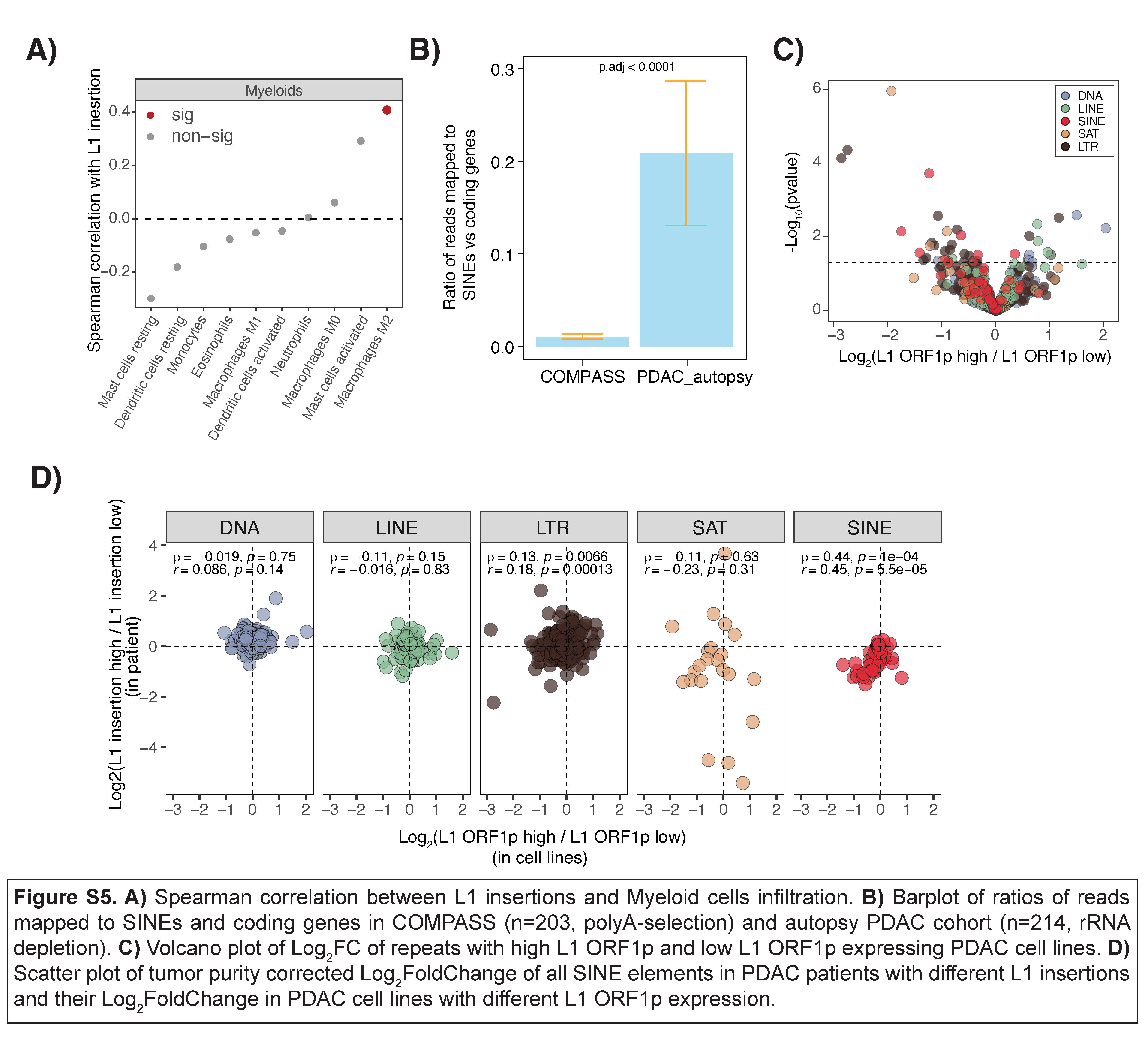

### Figure S6.png

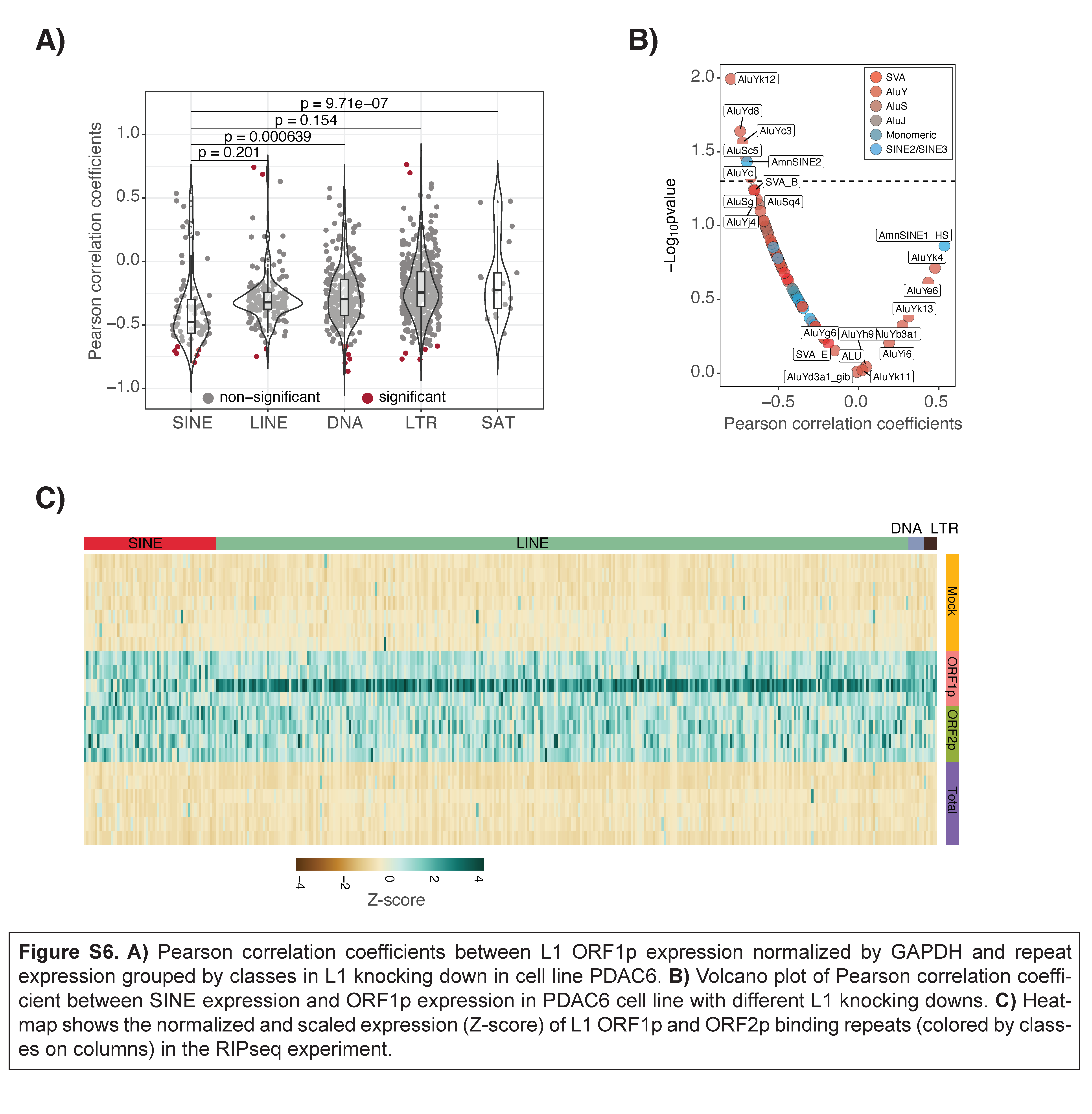

### Figure S7.png

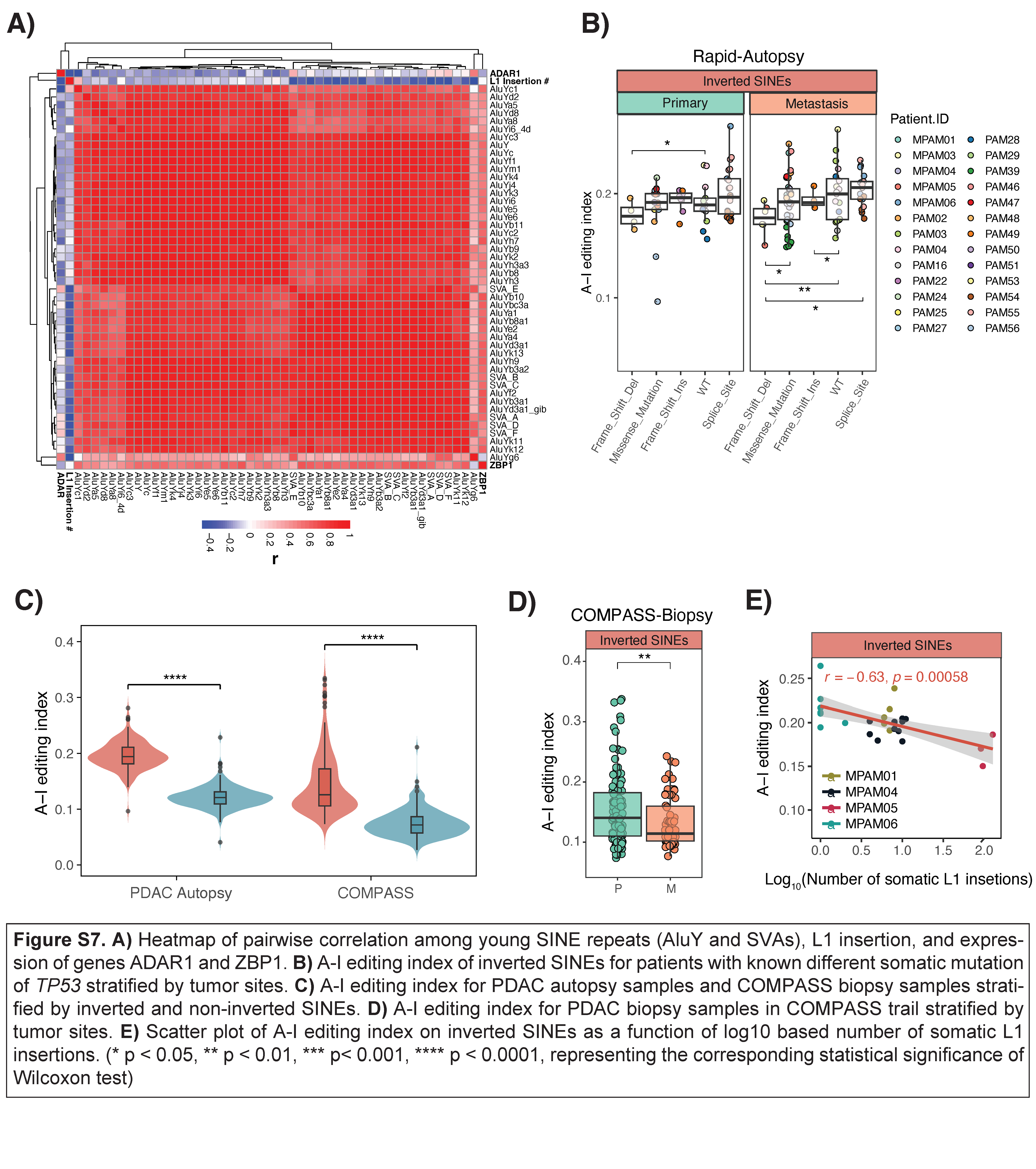
